# A Multi-omics Method Enabled by Sequential Metabolomics and Proteomics for Human Pluripotent Stem Cell-derived Cardiomyocytes

**DOI:** 10.1101/2021.06.23.449630

**Authors:** Elizabeth F. Bayne, Aaron D. Simmons, David S. Roberts, Yanlong Zhu, Timothy J. Aballo, Benjamin Wancewicz, Sean P. Palecek, Ying Ge

## Abstract

Human pluripotent stem cell-derived cardiomyocytes (hPSC-CMs) show immense promise for patient-specific disease modeling, cardiotoxicity screening, and regenerative therapy development. However, hPSC-CMs in culture have not recapitulated the structural or functional properties of adult CMs *in vivo* thus far. To gain global insight into hPSC-CM biology, we established a multi-omics method for analyzing the hPSC-CM metabolome and proteome from the same cell culture, creating multi-dimensional profiles of hPSC-CMs. Specifically, we developed a sequential extraction to capture metabolites and proteins from the same hPSC-CM monolayer cultures, and analyzed these extracts using high-resolution mass spectrometry (MS). Using this method, we annotated 205 metabolites/lipids and 4,008 proteins from 10^6^ cells with high reproducibility. We further integrated the proteome and metabolome measurements to create network profiles of molecular phenotypes for hPSC-CMs. Out of 310 pathways identified using metabolomics and proteomics, 40 pathways were considered significantly overrepresented (FDR-corrected *p* ≤ 0.05). Highly populated pathways included those involved in protein synthesis (ribosome, spliceosome), ATP generation (oxidative phosphorylation), and cardiac muscle contraction. This multi-omics method achieves deep coverage of metabolites and proteins, creating a multidimensional view of the hPSC-CM phenotype, which provides a strong technological foundation to advance the understanding of hPSC-CM biology.

## Introduction

Human pluripotent stem cell-derived cardiomyocytes (hPSC-CMs) have shown immense promise for patient-specific disease modeling, cardiotoxicity screening, and regenerative therapy development (1–5). Enabled by recent advances, hPSC-CMs can now be generated consistently with high yield and purity at a clinically relevant scale (6–9). However, one of the roadblocks impeding the realization of the full potential of hPSC-CMs is their structural and phenotypic immaturity compared to adult CMs (9, 10). Significant efforts have been made to mature hPSC-CMs and considerable progress has been achieved; nevertheless, hPSC-CMs still lack adult-like phenotypes *in vitro* (9, 11). As hPSC-CM maturation entails complex signaling networks, new discovery approaches are urgently needed to gain systems-level insights into hPSC-CM biology (9, 12, 13).

High-throughput “omics” technologies provide transformative insights to elucidate complex biological processes. Conceivably, a multi-omic strategy will offer a more comprehensive understanding of hPSC-CM complexity than any single omic technology alone (14–17). In the post-genomic era, proteomics and metabolomics offer an unbiased suite of tools to link proteins and metabolites for integrated analysis of signal transduction, cellular metabolism, and phenotype on a systems biology level (17–19). Analysis of lipids, metabolites, and proteins from the same sample can provide precise insights to the phenotype from which they arise, and elimination of parallel sample processing mitigates concerns around batch variability (15, 20). By measuring metabolites and proteins from the same hPSC-CM monolayer sample, the amount of information is maximized from these costly and time-consuming cultures. However, a multi-omics method with the capability to analyze both metabolome and proteome from the same hPSC-CM culture remains lacking. There are still challenges associated with reproducible sample extraction for both metabolomics and proteomics, as well as correlation of metabolite and protein measurements from the same cell culture.

Herein, we developed a novel multi-omics method enabled by a sequential extraction and high-resolution mass spectrometry (MS)-based metabolomics and proteomics to analyze monolayer cultures of hPSC-CMs. After acquisition of metabolite and protein measurements from the same cell culture, we create a multi-dimensional profile of molecular phenotypes of hPSC-CMs. This report represents the first integration of untargeted metabolomics and proteomics data from hPSC-CM monolayer cultures to identify prominent biological pathways of hPSC-CMs. We demonstrate high reproducibility in our metabolomics and proteomics measurements, which provides a strong technological foundation to better understand hPSC-CM biology.

### Experimental Procedures

#### hPSC and hPSC-CM Cell Culture and Validation

Human induced pluripotent stem cells (WTC11 cell line) were maintained and differentiated into cardiomyocytes using the GiWi protocol as described previously (21). hPSC-CM differentiation efficiency was assessed in parallel wells by flow cytometry for cardiac troponin T (cTnT)-positive CMs on Day 16 of differentiation (21). The resulting purity was 72 ± 2% cTnT+ (n=5) (**Figure S1**).

#### Sequential Extraction of hPSC-CM Culture

Metabolites and proteins were sequentially extracted using a solvent-based quench technique (22). Cells were incubated in 1.0 mL cold (4°C) 100% methanol for 1 minute to quench cellular metabolic activity. Subsequently, cells were scraped into a 1.5 mL microcentrifuge tubes in methanol, referred to as “Metabolite-Protein Mixture.” Metabolite-Protein Mixture was vortexed for 10s and incubated for 10 min at 4 °C. The Metabolite-Protein Mixture was centrifuged for 5 min at 14,000 g, creating a metabolite-rich supernatant and protein-rich cell pellet. The pellet was rinsed with 200 μL DPBS. Metabolite and protein samples were dried under vacuum and stored at −80 °C.

#### Untargeted Metabolomics of hPSC-CM

Metabolites were reconstituted in 1:1 methanol:water. FIE-FTICR MS-based metabolic fingerprinting was performed using a Waters ACQUITY UPLC M-Class System (Waters Corporation, Milford, MS, USA) coupled to a Bruker solariX 12T FTICR mass spectrometer (Bruker Daltonics, Bremen, Germany) without an LC column as described previously (23). MS data were processed using MetaboScape v2021 and feature assignments were manually validated using DataAnalysis v4.3 (both Bruker Daltonics, Bremen, Germany). Bucket lists for data collected in positive and negative ionization modes were generated using the T-Rex 2D algorithm and combined to create one merged bucket list using a mass error tolerance of < 1.0 ppm. Features were annotated by SmartFormula in MetaboScape as described previously (24) and by accurate mass using the Mass Bank of North America, Human Metabolome Database (HMDB) (25), LipidBlast (26), and METLIN databases (27) with an error tolerance ≤ 3.0 ppm. See **Supplemental File 1** for MetaboScape output. Overlap in metabolite features between hPSC-CMs was visualized using BioVenn (28). Metabolites were mapped to networks using MetScape plugin in CytoScape v 3.8.1 (29).

#### Global Proteomics Analysis of hPSC-CM

Proteins from the dried cell pellet were solubilized in 10 μL buffer containing 0.25% w/v Azo. Samples were normalized, reduced, alkylated, and digested with Trypsin Gold (Promega, Madison, Wisconsin, USA). Samples were quenched and irradiated at 305 nm for 5 min, desalted and dried under vacuum.

Nanoflow LC-MS analysis of tryptic peptides was performed using a timsTOF Pro (30) coupled to a nanoElute nano-flow UHPLC system using a Captivespray nano-electrospray ion source all Bruker Daltonics, Bremen, Germany). The timsTOF Pro was operated in Parallel Accumulation and Serial Fragmentation (PASEF) mode. Tandem mass spectra were searched against the UniProt human database UP000005640 using MaxQuant v1.6.17.0 (31) using a false discovery rate of 0.01. Label free quantification (LFQ) was performed using classic normalization and a minimum ratio count of 2. For MaxQuant output, refer to **Supplemental File 3**. To determine correlation of LFQ intensities across samples, Perseus software v1.6.14.0 (32) was used to transform the data by log2x and generate scatter plots with Pearson correlation coefficients. Overlap in protein groups and peptides identified between hPSC-CMs was visualized using BioVenn. Gene ontology and pathway analysis of proteins identified in the dataset was performed using the Database for Annotation, Visualization, and Integrated Discovery (DAVID) (33) .

#### Multi-omics (Metabolomics-Proteomics) Data Integration

Protein (UniProt identifications) and metabolite (KEGG identifiers) lists were uploaded to MetaboAnalyst 5.0 for integrative pathway analysis (34). The Joint Pathway Analysis module was used to determine significantly enriched pathways within the dataset (hypergeometric test, *p* ≤ 0.05).

For further details, refer to **Supplemental Materials and Methods.**

## Results and Discussion

### A Multi-Omics Platform

We have developed a novel multi-omics platform to analyze metabolites (including lipids) and proteins from the same hPSC-CM sample. To harvest metabolites and proteins, cellular activity was quenched using cold methanol (22), simultaneously extracting metabolites and precipitating proteins (**Figure 1**). This quenching protocol achieves rapid, reproducible access to the metabolome and proteome. After a brief centrifugation step, the metabolite-rich supernatant and the dehydrated cell pellet were easily separated and analyzed using MS-based metabolomics and proteomics, respectively. The supernatant required no further preparation prior to analysis by MS-based metabolomics, while the cell pellet was resolubilized using Azo before the analysis by the MS-based proteomics. Direct metabolite extraction using methanol was chosen over traditional cell-lifting based protocols as it has been shown to better preserve endogenous intracellular metabolite profiles (35).

**Figure 1.**
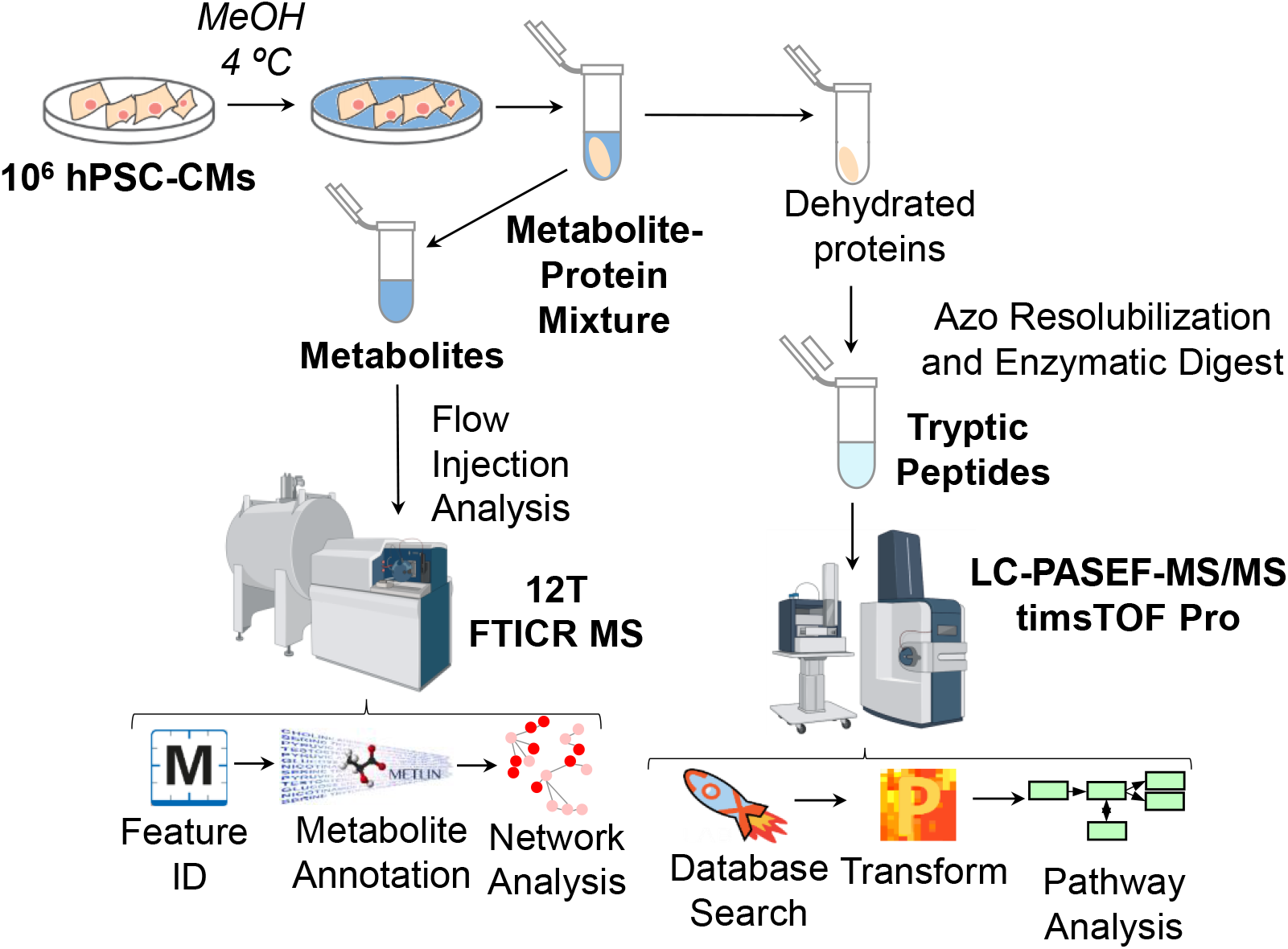
Schematic of sequential extraction of metabolites and proteins from hPSC-CM monolayer culture and MS analysis for integrated metabolomics and proteomics. Detailed extraction procedure and analysis is described in **Supplemental Materials and Methods**.

We used several specialized analytical approaches to enable a high throughput and comprehensive analysis of hPSC-CM metabolites and proteins. To measure metabolites, we used an automated flow injection electrospray (FIE) platform integrated to Fourier transform ion cyclotron resonance (FTICR) MS (24). The ultrahigh mass accuracy and resolving power of the FTICR MS enable simultaneous detection and quantification of complex mixture of metabolites without the need for chromatographic separation, sample fractionation, or chemical derivatization, resulting in metabolite identifications with sub-ppm mass accuracy and a high-throughput analytical workflow (24).

To solubilize proteins for global bottom-up proteomics, we used Azo (36), a novel photo-cleavable surfactant that enables high-throughput enzymatic digestion. Further, minimal sample cleanup is required as <5 minutes of exposure to UV light will degrade it into MS-compatible products (23). Moreover, Azo has been shown to effectively access critical membrane and ECM proteins (23, 37). To analyze peptides, we used online data-dependent Parallel Accumulation-Serial Fragmentation (dda-PASEF) on the timsTOF Pro (38, 39). The coupling of trapped ion mobility spectrometry (tims) with PASEF on the timsTOF Pro enables rapid sequencing of peptides, resulting in deep and sensitive proteome coverage using a standard 120 minute sample run (39).

### High Resolution Untargeted Metabolomics enabled by FIE-FTICR MS Platform

Metabolite samples were directly infused into a 12 T FTICR MS using an automated flow injection and electrospray ionization (FIE-FTICR MS) (**Figure 2A**). Our FIE-FTICR MS platform requires only 5 min of analysis time per sample and detects hundreds of metabolite features with high mass accuracy (**Figure S2A**). Additionally, the FIE-FTICR MS platform can profile small molecule metabolites as well as lipid composition of the sample within this 5 min analysis. FIE-FTICR MS enables fast polarity switching, allowing for deeper coverage of unique metabolites in the sample, including amino acids, alkenes, and ceramides in the positive mode and a variety of phospholipid classes in the negative ionization mode (**Figure S2B**). From approximately 10^6^ cells, we detected an average of 626 metabolite features between positive and negative ionization modes. Of those features, 468 were annotated by the SmartFormula (SF) function in MetaboScape with ≤ 5.0 ppm mass error tolerance, generating a putative chemical formula based exact mass and isotopic pattern. In a representative zoomed-in window of the mass spectrum ranging from 180-250 *m/z* showed 14 SF annotations made within 5 ppm error in positive ionization mode, with 12 features annotated with errors around 2 ppm (**Figure S3A**). In the negative ionization mode, 12 features were annotated by SF in the same 70 *m/z* window within 5 ppm error, with 7 of them annotated with an *m/z* error tolerance of < 1 ppm (**Figure S3B**). Metabolites were annotated from databases such as METLIN and Mass Bank of North America were annotated using high mass accuracy (≤ 3.0 ppm). These annotations cover a broad range of metabolites and lipids, including triglycerides (TAG), ATP, NADH, and UDP-glucose (**Figure 2A, Supplemental File 2**).

**Figure 2.**
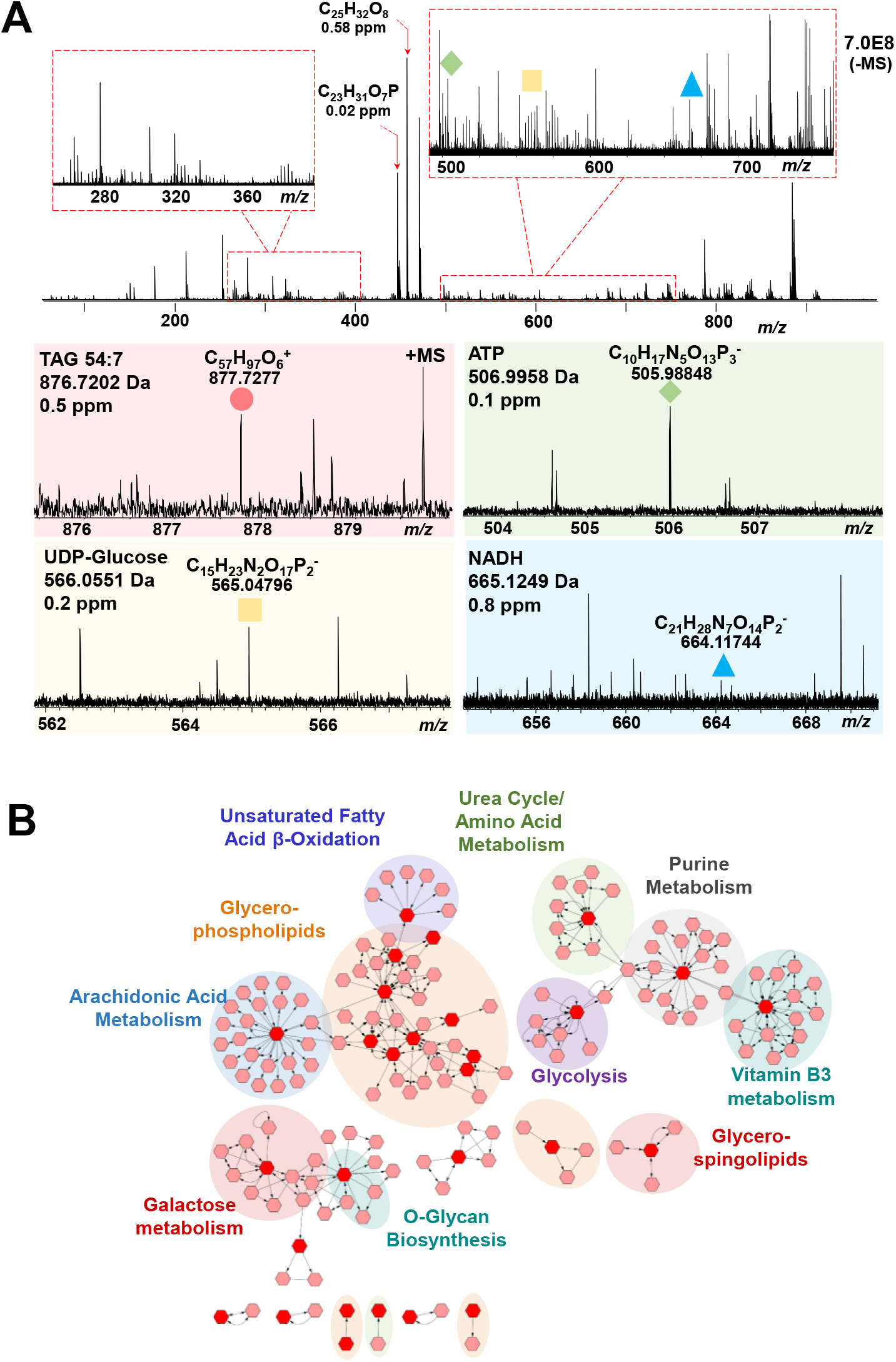
Ultrahigh-resolution mass spectrometry (MS)-based metabolomics of hPSC-CM cell culture. **A)** Identification of metabolites from complex broadband spectra enabled by FIE-FTICR MS. Important metabolites are identified with sub-ppm mass accuracy, including stearic acid (upper left), NADH (upper right), UDP-glucose (lower left), and cardiolipin (lower right). **B)** Network analysis of identified metabolites show arachidonic acid metabolism, glycolysis, and glycerophospholipid metabolism pathways represented in the dataset.

We demonstrated highly reproducible injections for the analysis of hPSC-CM metabolites (**Figure S4A**). Mass spectra of triplicate injections show similarity in metabolite feature intensity and populations in both positive and negative ionization modes. After a blank reduction step, 624-629 features were detected in each injection replicate and shared 623 features in common, corresponding to over 99% overlap between triplicate injections (**Figure S4B**). Variation in shared feature intensities was low, with 6.9% median relative standard deviation (RSD) for all features detected. 96% of features detected had a signal intensity variation less than 20% RSD and 72% of features less than 10% RSD (**Figure S4C**). We used this strong technical foundation to evaluate the reproducibility of hPSC-CM metabolite extraction using biological replicates (n=3). Visual comparison of mass spectra shows similar intensities and feature population between hPSC-CM biological replicates in both ionization modes (**Figure S5A**). 619-623 features were detected in 3 of 3 triplicate injections for each biological replicate and 613 features were shared between them, corresponding to 98% overlap of features detected between biological replicates (**Figure S5B**). Of these shared features, 459 were annotated with SF. Variation in feature intensity across biological replicates was 13% median RSD, and 84% of features varied by ≤ 20% RSD (**Figure S5C**). Taken with the high overlap in features detected between biological replicates and high technical reproducibility, these variation in the data indicate the methods’ sensitivity to differences in metabolite profiles between hPSC-CMs.

Hundreds of metabolites important to hPSC-CM biology were identified in our dataset with high mass accuracy, including those involved in pathways such as amino acid metabolism, glycolysis, and glycerophospholipid metabolism (**Figure 2B**). 205 metabolites were annotated using a combination of the METLIN, HMDB, LipidBlast, and MoNA databases with an error tolerance of ≤ 3.0 ppm. Of those metabolites, 46 were annotated with mass error tolerance of ≤ 0.2 ppm, and 117 were annotated ≤ 1.0 ppm. Energy-yielding substrates such as glucose, ATP, fatty acids, acyl-carnitines, amino acids, and triglyceride (TAG) species were detected in the dataset (**Table S1, Supplemental File 2**), which can be correlated to energy-generating pathways. Metabolites involved in oxidative phosphorylation (e.g. NAD+/NADH, ATP/ADP), a primary source for ATP generation in mature CMs (40, 41), were also consistently identified in the dataset. Diverse phospholipid species including sphingolipids, ceramides, phosphocholines (PC), cardiolipins (CL), phosphotidylserines (PS), phosphotidylglycerols (PG), and phosphotidylethanolamines (PE) were also identified in the dataset (**Table S2**). Phospholipids have been shown to play a key role in the position and function of membrane-embedded ion transporter proteins, which regulate cardiac contraction (42). Thus, we can access a wide range of biologically relevant metabolite classes, from carbohydrates to lipids, using this fast and reproducible procedure to investigate active metabolic pathways in hPSC-CM biology.

### Global proteomics of the hPSC-CM cell pellet

After metabolite extraction, we subsequently extracted proteins from the hPSC-CM cell pellet and analyzed the resulting peptides using label-free global bottom-up proteomics. The pellet was resolubilized in a buffer containing MS-compatible surfactant Azo (23), enzymatically digested, and analyzed by online LC-PASEF-MS using the timsTOF Pro (**Figure 1**) (39). Extraction reproducibility was demonstrated for the Azo-enabled bottom-up global proteomics method through similar protein yields for each pellet as demonstrated by Bradford assay, with protein yields of 26-31 μg of total protein per hPSC-CM sample (**Tables S3, S4**). Given that only 100-200 ng peptides are required for each bottom-up global proteomics run, this procedure yields adequate protein to analyze each hPSC-CM monolayer culture and could be scaled down to accommodate smaller monolayer cultures (<10^6^ cells). The online data-dependent PASEF proteomics platform showed high technical reproducibility and robustness throughout the analysis. Over the course of 29 sample injections (58 hours, 2.5 days), the maximum shift in retention time is only 0.2 min difference between injections of the K562 whole cell lysate standard at 200 ng at the beginning and end of the run, demonstrating impressive run-to-run reproducibility across multiple samples injections (**Figure S6A**). Additionally, label-free quantitation (LFQ) of the K562 cell standard showed strong correlation between three bracketing standard runs spanning 58 hours, with a Pearson correlation coefficient of 0.997 (**Figure S6B**).

Global proteomics revealed a large diversity in protein groups recovered from the cell pellet after the initial metabolite extraction. Analysis of online dda-PASEF bottom-up proteomics data using a false discovery rate (FDR) of 1% at the protein and peptide level revealed a total of 31,617 unique peptides between three hPSC-CM biological replicates, corresponding to 4,319 combined protein groups. Using only 200 ng per run, 21,240 peptides were shared between three hPSC-CM cultures. We found 3,663 proteins in common between replicates, corresponding to 85% overlap between biological replicates (**Figure 3A**). Additionally, strong correlation of LFQ intensities show consistent recovery of proteins from the dual extraction, with Pearson correlation coefficients ranging from 0.960-0.983 (**Figure 3B**). Normal unimodal distribution of log transformed (LFQ) intensities was observed for each of the three replicates. These results illustrate the reproducible extractions of proteins from the residual hPSC-CM pellet by this sequential extraction strategy and demonstrate the feasibility of applying this method for quantitation between samples from sequentially-extracted hPSC-CMs for metabolomics and proteomics.

**Figure 2.**
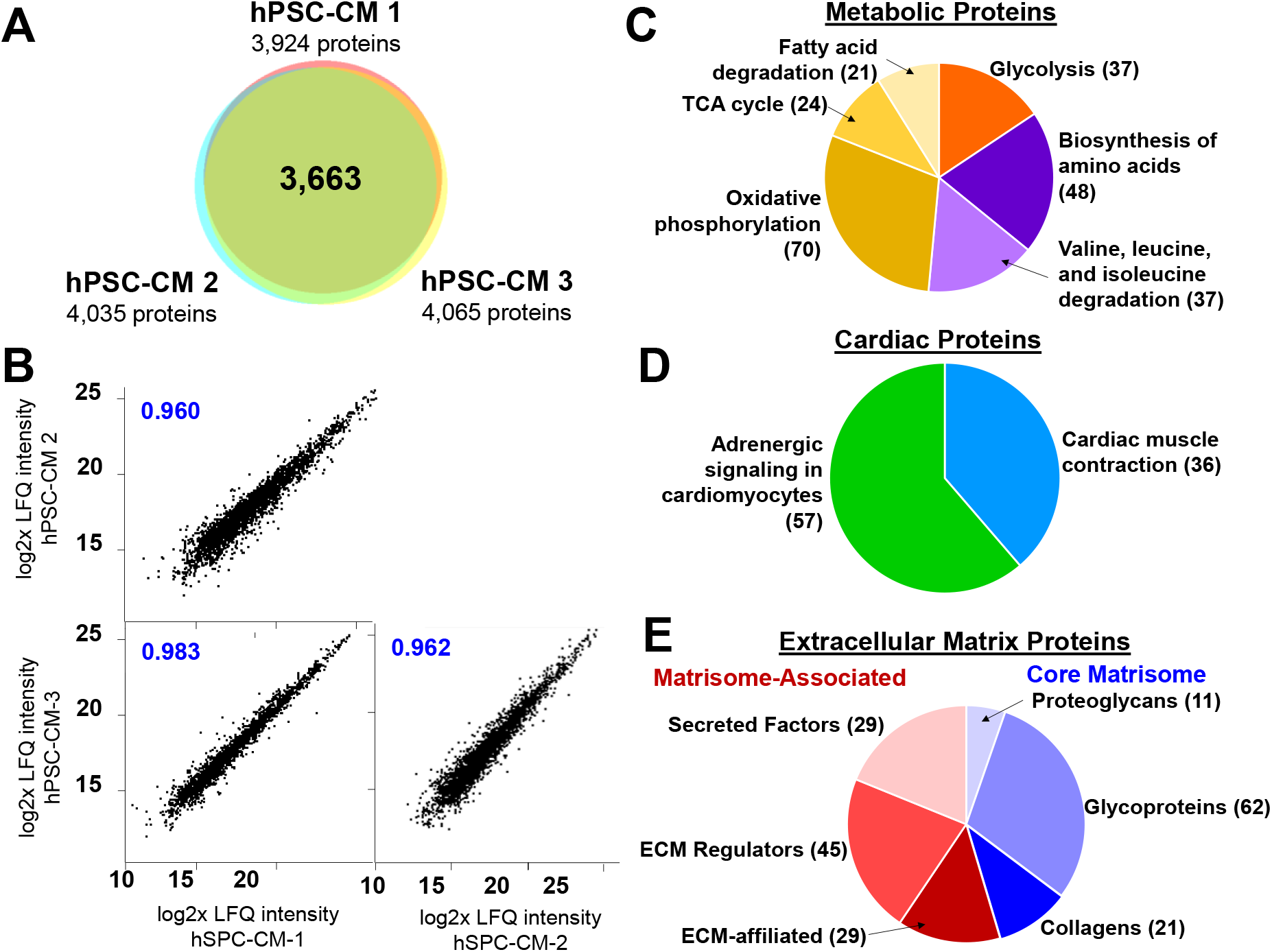
Global proteomics of hPSC-CM cell pellet and metabolomics and pathways revealed by integrated multi-omics analysis. **A)** Reproducibility in hPSC-CM biological replicates (n=3). Protein groups identified show 85% overlap between three hPSC-CM monolayer cultures, corresponding to 3,663 protein groups in common. **B)** Log2x-transformed LFQ intensities of shared proteins in each biological replicate show strong correlation between samples, with Pearson correlation coefficients ranging from 0.960-0.983. **C)** Representative KEGG pathways considered statistically-overrepresented (FDR-corrected *p* ≤ 0.05) for Metabolic Proteins **(C)** and Cardiac-specific proteins (**D**). Metabolic proteins included those involved in glycolysis (hsa00010, orange), amino acid metabolism (biosynthesis of amino acids, hsa01230, and valine, leucine, and isoleucine degradation, hsa00280, both purple), and oxidative respiration (oxidative phosphorylation, hsa00190, TCA cycle, hsa00020, and fatty acid degradation, hsa00071, all yellow). Proteins involved in cardiac-specific pathways such as adrenergic signaling of cardiomyocytes (hsa04261, green) and cardiac muscle contraction (hsa04260, blue) were identified. The number of proteins in each pathway is indicated in parentheses. **E)** Extracellular matrix (ECM) proteins were categorized into Core Matrisome (red) and Matrisome-associated proteins (blue). The number of proteins in each category is indicated in parentheses. Core matrisome proteins include proteoglycans, glycoproteins, and collagen species, while matrisome-associated proteins include secreted factors, ECM regulators, and ECM-affiliated proteins.

Gene ontology analysis revealed highly diverse protein groups recovered from the sequentially-extracted hPSC-CM pellet. Protein groups within our dataset were represented by KEGG pathways using the Database for Annotation, Visualization, and Integrated Discovery (DAVID) (33). Using this tool, we found 91 KEGG pathways represented in the proteomics dataset. Among them, 65 pathways were considered abundantly represented (FDR-corrected *p* ≤ 0.05), many of which reflect ATP-generating pathways as well as cardiomyocyte function and development (**Table S5, Supplemental File 4**). For example, two of the most significantly abundant pathways represented in the dataset, the Spliceosome and Ribosome, are relevant to protein processing and translation.

Proteins involved in metabolic pathways were highly represented throughout the dataset, constituting 417 total protein groups (**Figure 3C**). Among them, diverse pathways were identified, such as amino acid metabolism pathways and mitochondrial fatty acid β-oxidation. Glycolysis is responsible for the majority of ATP generation in the immature hPSC-CMs (40), and was represented by 37 proteins in the dataset, including hexokinase-1 (HK-1, gene: *HK1*), α-enolase (gene: *ENO1*), and pyruvate kinase (gene: *PKM*). Fatty acid degradation comprised 24 proteins in the dataset, including mitochondrial proteins glutaryl-CoA dehydrogenase (gene: *GCDH*) and very long-chain specific acyl-CoA dehydrogenase (gene: *ACADVL*). Metabolism of amino acids and derivatives, including valine, leucine, and isoleucine degradation (hsa00280) and biosynthesis of amino acids (hsa01230) was represented by 39 and 48 proteins, respectively. The depth of coverage of metabolic protein groups and pathways in the hPSC-CM dataset can be used for metabolite annotations to analyze the cells from a multi-omics perspective.

Cardiac-specific proteins were also consistently detected throughout the dataset. Cardiac-specific pathways found in KEGG pathways included Cardiac Muscle Contraction (hsa02460) and Adrenergic Signaling in Cardiomyocytes (hsa02461) comprising 36 and 57 proteins, respectively (**Figure 3D**). Cardiomyocyte-specific marker cardiac troponin T (cTnT, gene: *TNNT2*) (8) was consistently detected in each CM replicate with 0.2% RSD in LFQ intensity. Other cardiac proteins such as slow skeletal troponin I (ssTnI, gene: *TNNI1*), α-cardiac muscle actin (α-CAA, gene: *ACTC1*), cardiac troponin C (cTnC, gene: *TNNC1*), cardiac myosin binding protein C (gene: *MYBPC3*), cardiac phospholamban (gene: *PLN*), and cardiomyocyte transcription factor GATA-4 (gene: *GATA4*) (43) were consistently detected as well (**Figure S7**). Important cardiac contractile proteins and markers of CM specification were consistently identified, including the atrial and ventricular isoforms of myosin light chain (MLC-2a, gene: *MYL7* and MLC-2v, gene: *MYL2*, respectively), and α- and β-myosin heavy chains (α-MHC, gene: *MYH6*, β-MHC, gene: *MYH7*, respectively). MLC-2v and MLC-2a are considered markers of development and the presence of MLC-2v can also indicate development of ventricular CM linages (10).

Importantly, a large number of proteins associated with the extracellular matrix (ECM) sub-proteome (“matrisome”) were detected in the sequentially-extracted hPSC-CMs (**Figure 3E**). Cardiac ECM proteins are crucial to providing structural support to the CM monolayer and help to withstand mechanical stress from continual contraction of the sarcomere (44). Key ECM proteins to CM function (44) were detected in the sequentially-extracted cultures, including collagen I (genes: *COL1A1*, *COL1A2*), collagen III (gene: *COL3A1*), and basement membrane proteins collagen IV (genes: *COL4A1, COL4A2, COL4A3* and *COL4A6*) and fibronectin (gene: *FN1*). Overall, core matrisome proteins (45) were consistently detected throughout the dataset, including 21 collagen species, 62 glycoproteins, and 11 proteoglycans. We also detected 115 matrisome-associated proteins, which are ECM regulators and secreted factors, or are structurally affiliated with the ECM (**Supplemental File 1**) (45). 43 proteins were associated with the KEGG pathway “ECM-receptor interaction” (hsa04512) including various subunits of the transmembrane integrin receptor complex (genes: *ITGB1* and *3*, *ITGAV*, and *ITGA1, 2*, and *5-7*), as well as α-, β-, and γ- chains of the laminin complex (genes: *LAMA1-LAMA5*, *LAMB1* and *2*, and *LAMC1* and *2*). The reliable detection of diverse matrisome species using a sequentially-extracted method is significant because protein solubility is the primary challenge of ECM proteomics (46, 47). Often, multi-step extraction procedures are used to isolate the ECM. Using a single-step proteomics extraction enabled by Azo, we detected a diverse array of core and associated matrisome constituents in addition to the metabolic and sarcomeric subproteomes, enabling analysis of each subproteome on a global scale.

### Integrative Multi-Omics Reveals Highly Populated Biological Pathways

Finally, we integrated the metabolomics and proteomics results to reveal pathways abundantly represented in the data using the Joint Pathway Analysis module in MetaboAnalyst (34). Here, metabolites and proteins were mapped to KEGG pathways. Out of 310 identified pathways, 25 were considered statistically enriched by hypergeometric test (FDR-corrected *p* ≤ 0.05), with many of them corresponding to metabolic processes within the KEGG global metabolic network, including fatty acid metabolism, pentose phosphate pathway, glycolysis and TCA cycle, purine metabolism, and amino acid metabolism (**Figure 4A, Supplemental File 5**). Metabolite and protein constituents of each pathway were identified using MetaboAnalyst and were mapped for further analysis. To look closer at the identified pathway results, we used the Joint Pathway Analysis module to visualize 55 proteins and 3 general metabolite classes (“branched fatty acids”) represented in the fatty acid biosynthesis and degradation pathways (**Figure 4B**). Other metabolic pathways abundantly represented in the dataset included glycerophospholipid metabolism, oxidative phosphorylation, and cell junction, (i.e. focal adhesion, tight junctions). Combined, 84 metabolite and protein hits were mapped to the oxidative phosphorylation pathway, including energy-generating metabolites NAD+/NADH and members of the electron transport chain and TCA cycle (**Figure 5**). Oxidative metabolism generates the majority of ATP in mature CM phenotypes (41), and mitochondrial pathways TCA cycle and oxidative phosphorylation comprised 24 and 74 proteins in the dataset, respectively, including each constituent of the membrane-embedded electron transport chain. Pathways involved in protein synthesis such as RNA transport and degradation, and the spliceosome were also abundantly represented (**Figure S8; Table S6**). Together, these results offer holistic insights into which cellular pathways are most active in d42 hPSC-CMs.

**Figure 4.**
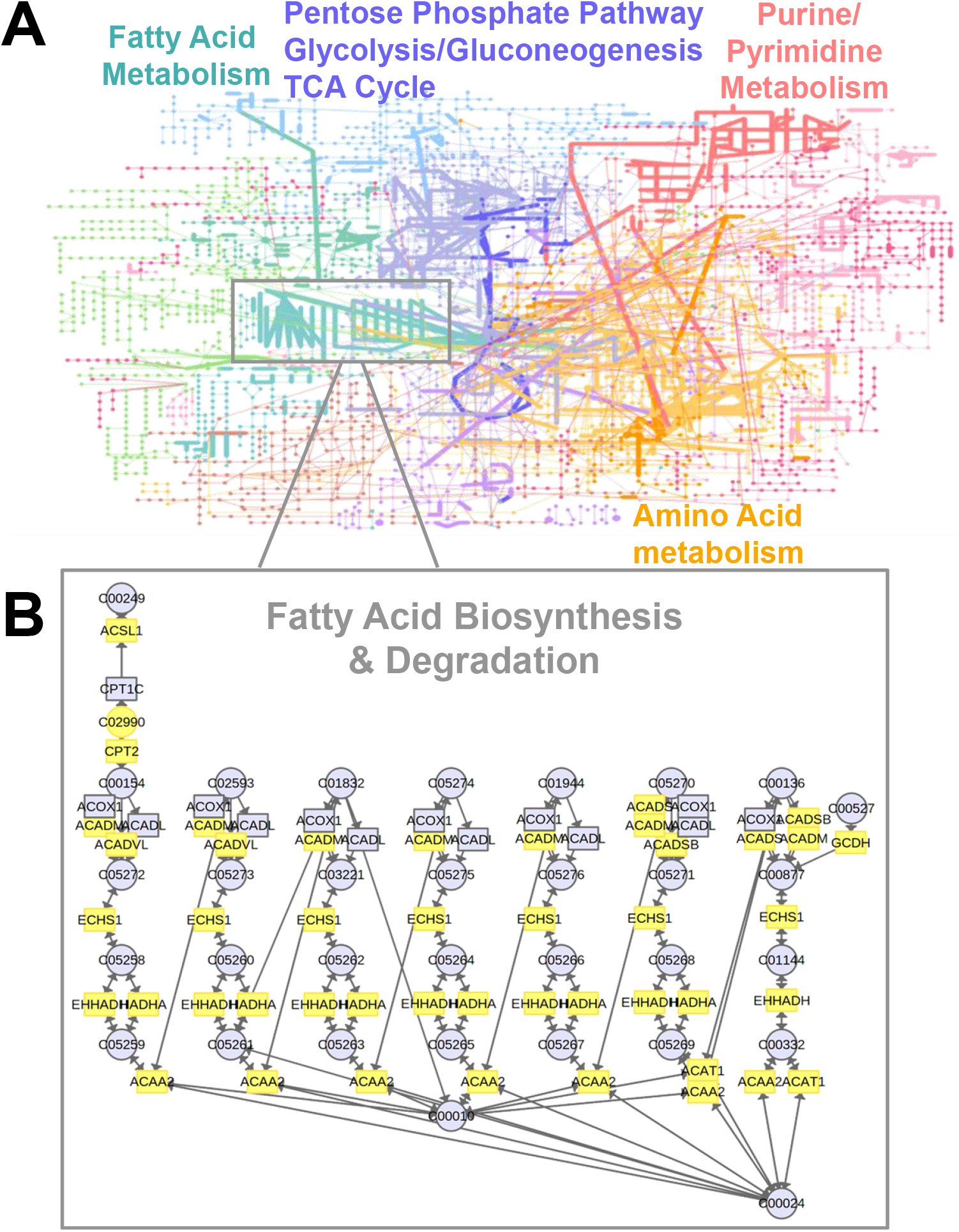
Integrative Multi-Omics Reveals Highly-Populated Biological Pathways. **A)** Visualization of multi-omics metabolism pathways by MetaboAnalyst. Proteomics and metabolomics data were uploaded to MetaboAnalyst for pathway analysis. The Network Analysis module showed significantly-enriched pathways within the KEGG metabolic network for proteins and metabolites, including fatty acid metabolism, amino acid metabolism, purine/pyrimidine metabolism, and glycolysis, pentose phosphate pathway, and TCA cycle. **B)** Zoom panel shows detailed pathways showing detected entities for fatty acid degradation. In total, 58 members of fatty acid degradation pathway were detected, indicated in yellow. KEGG nomenclature is used to describe proteins and metabolites.

**Figure 5.**
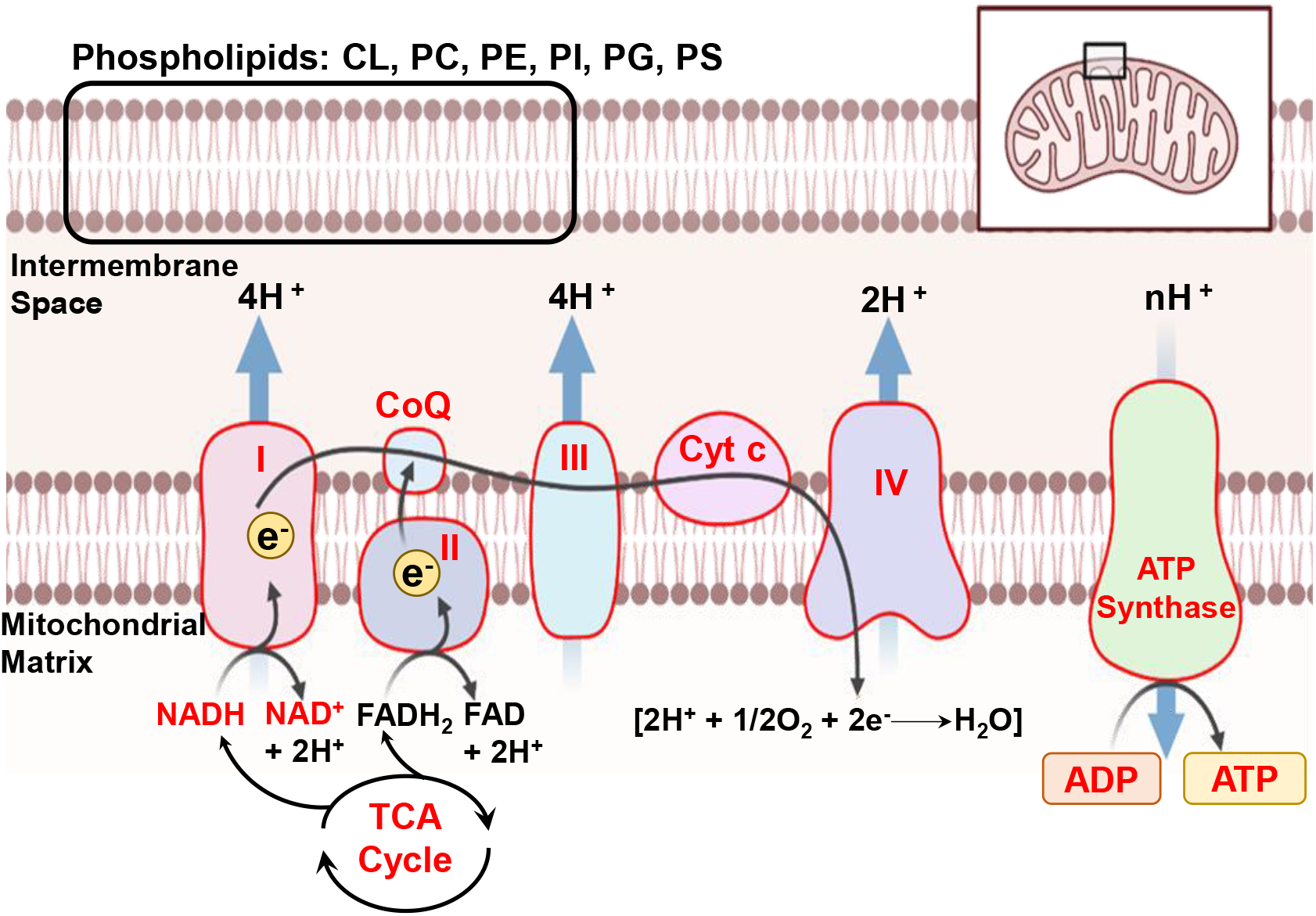
Prominent energy-generating pathways in hPSC-CMs revealed by protein and metabolites. KEGG pathway oxidative phosphorylation was represented by a total of 84 protein and metabolite hits within the dataset, highlighted in red. Additionally, many phospholipid species that comprise the mitochondrial membrane were detected, including cardiolipins (CL), phosphocholines (PC), phosphoethanolamines (PE), phosphoinositols (PI), phosphoglycerols (PG), and phosphatidylserines (PS).

## Conclusions

Our novel multi-omics method for the analysis of metabolites and proteins from the same hPSC-CM monolayer culture enables an integrated assessment of the hPSC-CM proteome and metabolome comprehensively and reproducibly. Benefiting from the rapid metabolite extraction from hPSC-CM cell culture and the facile sample processing for MS analysis, a large number of samples can be extracted and analyzed. We achieved broad coverage of metabolite classes from a rapid methanol extraction of hPSC-CM in culture, including carbohydrates, amino acids, and a variety of phospholipids. Strong technical and biological reproducibility was shown by highly similar mass spectra and metabolite annotations. Additionally, we achieved high proteome coverage of the residual hPSC-CM cell pellet, identifying approximately 4,000 proteins per hPSC-CM pellet. We also showed consistent recovery of protein from each hPSC-CM monolayer. Using 200 ng per sample, we measured strong correlation of protein intensities from sequentially-extracted cultures, and identified a wide range variation of proteins, spanning metabolic, sarcomeric, and extracellular matrix proteins. Finally, we mapped identified metabolites and proteins to networks and identified highly abundant pathways in the dataset. We envision this multi-omics strategy will provide a technical foundation for future studies on the quantitative evaluation of various hPSC-CM conditions and identification of integrated proteomic and metabolic markers to advance our understanding of hPSC-CM biology.

## Acknowledgements

The financial support is provided by NIH grant NIH R01HL148059 (to E.F.B.), NIH 5 T32 GM135066 (to A.D.S.), and NSF grant EEC-1648035 (to S.P.P.). D.S.R. would like to acknowledge support from the American Heart Association Predoctoral Fellowship Grant No. 832615 / David S. Roberts / 2021. B.W. acknowledges support from the NIH Molecular and Pathology Training Program T32GM081061 and F31HL152647 (to B.W.). Y. G. would like to acknowledge NIH R01HL096971, R01GM125085, and S10OD018475. We thank Julia Zhou for assistance with metabolite database searching.

## Author Contributions

E.F.B., A.D.S., D.S.R., Y.Z., and T.J.A. performed the research; E.F.B., A.D.S., D.S.R., Y.Z., T.J.A., B.W., S.P.P. and Y.G. wrote the manuscript.

## Notes

### Competing Interest Statement

The authors have declared no competing interest.

## References

1. Zhang, J.; Wilson Gisela, F.; Soerens Andrew, G.; Koonce Chad, H.; Yu, J.; Palecek Sean, P.; Thomson James, A.; Kamp Timothy, J., Functional Cardiomyocytes Derived From Human Induced Pluripotent Stem Cells. Circulation Research 2009, 104, (4), e30–e41.

2. Burridge, P. W.; Li, Y. F.; Matsa, E.; Wu, H.; Ong, S.-G.; Sharma, A.; Holmström, A.; Chang, A. C.; Coronado, M. J.; Ebert, A. D.; Knowles, J. W.; Telli, M. L.; Witteles, R. M.; Blau, H. M.; Bernstein, D.; Altman, R. B.; Wu, J. C., Human induced pluripotent stem cell–derived cardiomyocytes recapitulate the predilection of breast cancer patients to doxorubicin-induced cardiotoxicity. Nature Medicine 2016, 22, (5), 547–556.

3. Ye, L.; Chang, Y.-H.; Xiong, Q.; Zhang, P.; Zhang, L.; Somasundaram, P.; Lepley, M.; Swingen, C.; Su, L.; Wendel, Jacqueline S.; Guo, J.; Jang, A.; Rosenbush, D.; Greder, L.; Dutton, James R.; Zhang, J.; Kamp, Timothy J.; Kaufman, Dan S.; Ge, Y.; Zhang, J., Cardiac Repair in a Porcine Model of Acute Myocardial Infarction with Human Induced Pluripotent Stem Cell-Derived Cardiovascular Cells. Cell Stem Cell 2014, 15, (6), 750–761.

4. Matsa, E.; Burridge, P. W.; Wu, J. C., Human stem cells for modeling heart disease and for drug discovery. Science translational medicine 2014, 6, (239), 239ps6–239ps6.

5. van Meer, B. J.; Tertoolen, L. G. J.; Mummery, C. L., Concise Review: Measuring Physiological Responses of Human Pluripotent Stem Cell Derived Cardiomyocytes to Drugs and Disease. Stem cells (Dayton, Ohio) 2016, 34, (8), 2008–2015.

6. Lian, X.; Hsiao, C.; Wilson, G.; Zhu, K.; Hazeltine, L. B.; Azarin, S. M.; Raval, K. K.; Zhang, J.; Kamp, T. J.; Palecek, S. P., Robust cardiomyocyte differentiation from human pluripotent stem cells via temporal modulation of canonical Wnt signaling. Proceedings of the National Academy of Sciences 2012, 109, (27), E1848.

7. Burridge, P. W.; Matsa, E.; Shukla, P.; Lin, Z. C.; Churko, J. M.; Ebert, A. D.; Lan, F.; Diecke, S.; Huber, B.; Mordwinkin, N. M.; Plews, J. R.; Abilez, O. J.; Cui, B.; Gold, J. D.; Wu, J. C., Chemically defined generation of human cardiomyocytes. Nature Methods 2014, 11, (8), 855–860.

8. Waas, M.; Weerasekera, R.; Kropp, E. M.; Romero-Tejeda, M.; Poon, E. N.; Boheler, K. R.; Burridge, P. W.; Gundry, R. L., Are These Cardiomyocytes? Protocol Development Reveals Impact of Sample Preparation on the Accuracy of Identifying Cardiomyocytes by Flow Cytometry. Stem Cell Reports 2019, 12, (2), 395–410.

9. Karbassi, E.; Fenix, A.; Marchiano, S.; Muraoka, N.; Nakamura, K.; Yang, X.; Murry, C. E., Cardiomyocyte maturation: advances in knowledge and implications for regenerative medicine. Nature Reviews Cardiology 2020, 17, (6), 341–359.

10. Feric, N. T.; Radisic, M., Maturing human pluripotent stem cell-derived cardiomyocytes in human engineered cardiac tissues. Advanced Drug Delivery Reviews 2016, 96, 110–134.

11. Ronaldson-Bouchard, K.; Ma, S. P.; Yeager, K.; Chen, T.; Song, L.; Sirabella, D.; Morikawa, K.; Teles, D.; Yazawa, M.; Vunjak-Novakovic, G., Advanced maturation of human cardiac tissue grown from pluripotent stem cells. Nature 2018, 556, (7700), 239–243.

12. Cai, W.; Zhang, J.; de Lange, W. J.; Gregorich, Z. R.; Karp, H.; Farrell, E. T.; Mitchell, S. D.; Tucholski, T.; Lin, Z.; Biermann, M.; McIlwain, S. J.; Ralphe, J. C.; Kamp, T. J.; Ge, Y., An Unbiased Proteomics Method to Assess the Maturation of Human Pluripotent Stem Cell–Derived Cardiomyocytes. Circulation Research 2019, 125, 936–953.

13. Melby, J. A.; de Lange, W. J.; Zhang, J.; Roberts, D. S.; Mitchell, S. D.; Tucholski, T.; Kim, G.; Kyrvasilis, A.; McIlwain, S. J.; Kamp, T. J.; Ralphe, J. C.; Ge, Y., Functionally Integrated Top-Down Proteomics for Standardized Assessment of Human Induced Pluripotent Stem Cell-Derived Engineered Cardiac Tissues. Journal of Proteome Research 2021, 20, (2), 1424–1433.

14. Nakayasu, E. S.; Nicora, C. D.; Sims, A. C.; Burnum-Johnson, K. E.; Kim, Y.-M.; Kyle, J. E.; Matzke, M. M.; Shukla, A. K.; Chu, R. K.; Schepmoes, A. A.; Jacobs, J. M.; Baric, R. S.; Webb-Robertson, B.-J.; Smith, R. D.; Metz, T. O., MPLEx: a Robust and Universal Protocol for Single-Sample Integrative Proteomic, Metabolomic, and Lipidomic Analyses. mSystems 2016, 1, (3), e00043–16.

15. Coman, C.; Solari, F. A.; Hentschel, A.; Sickmann, A.; Zahedi, R. P.; Ahrends, R., Simultaneous Metabolite, Protein, Lipid Extraction (SIMPLEX): A Combinatorial Multimolecular Omics Approach for Systems Biology. Molecular & cellular proteomics : MCP 2016, 15, (4), 1453–1466.

16. Karczewski, K. J.; Snyder, M. P., Integrative omics for health and disease. Nature Reviews Genetics 2018, 19, (5), 299–310.

17. Joshi, A.; Rienks, M.; Theofilatos, K.; Mayr, M., Systems biology in cardiovascular disease: a multiomics approach. Nature Reviews Cardiology 2020.

18. Mayr, M.; Madhu, B.; Xu, Q., Proteomics and Metabolomics Combined in Cardiovascular Research. Trends in Cardiovascular Medicine 2007, 17, (2), 43–48.

19. McGarrah Robert, W.; Crown Scott, B.; Zhang, G.-F.; Shah Svati, H.; Newgard Christopher, B., Cardiovascular Metabolomics. Circulation Research 2018, 122, (9), 1238–1258.

20. Pinu, F. R.; Beale, D. J.; Paten, A. M.; Kouremenos, K.; Swarup, S.; Schirra, H. J.; Wishart, D., Systems Biology and Multi-Omics Integration: Viewpoints from the Metabolomics Research Community. Metabolites 2019, 9, (4).

21. Lian, X.; Zhang, J.; Azarin, S. M.; Zhu, K.; Hazeltine, L. B.; Bao, X.; Hsiao, C.; Kamp, T. J.; Palecek, S. P., Directed cardiomyocyte differentiation from human pluripotent stem cells by modulating Wnt/β-catenin signaling under fully defined conditions. Nature Protocols 2013, 8, (1), 162–175.

22. Teng, Q.; Huang, W.; Collette, T. W.; Ekman, D. R.; Tan, C., A direct cell quenching method for cell-culture based metabolomics. Metabolomics 2008, 5, (2), 199.

23. Brown, K. A.; Tucholski, T.; Eken, C.; Knott, S.; Zhu, Y.; Jin, S.; Ge, Y., High-Throughput Proteomics Enabled by a Photocleavable Surfactant. Angewandte Chemie International Edition 2020, 59, (22), 8406–8410.

24. Zhu, Y.; Wancewicz, B.; Schaid, M.; Tiambeng, T. N.; Wenger, K.; Jin, Y.; Heyman, H.; Thompson, C. J.; Barsch, A.; Cox, E. D.; Davis, D. B.; Brasier, A. R.; Kimple, M. E.; Ge, Y., Ultrahigh-Resolution Mass Spectrometry-Based Platform for Plasma Metabolomics Applied to Type 2 Diabetes Research. Journal of Proteome Research 2021, 20, (1), 463–473.

25. Wishart, D. S.; Feunang, Y. D.; Marcu, A.; Guo, A. C.; Liang, K.; Vázquez-Fresno, R.; Sajed, T.; Johnson, D.; Li, C.; Karu, N.; Sayeeda, Z.; Lo, E.; Assempour, N.; Berjanskii, M.; Singhal, S.; Arndt, D.; Liang, Y.; Badran, H.; Grant, J.; Serra-Cayuela, A.; Liu, Y.; Mandal, R.; Neveu, V.; Pon, A.; Knox, C.; Wilson, M.; Manach, C.; Scalbert, A., HMDB 4.0: the human metabolome database for 2018. Nucleic Acids Research 2018, 46, (D1), D608–D617.

26. Kind, T.; Liu, K.-H.; Lee, D. Y.; DeFelice, B.; Meissen, J. K.; Fiehn, O., LipidBlast in silico tandem mass spectrometry database for lipid identification. Nature Methods 2013, 10, (8), 755–758.

27. Guijas, C.; Montenegro-Burke, J. R.; Domingo-Almenara, X.; Palermo, A.; Warth, B.; Hermann, G.; Koellensperger, G.; Huan, T.; Uritboonthai, W.; Aisporna, A. E.; Wolan, D. W.; Spilker, M. E.; Benton, H. P.; Siuzdak, G., METLIN: A Technology Platform for Identifying Knowns and Unknowns. Analytical Chemistry 2018, 90, (5), 3156–3164.

28. Hulsen, T.; de Vlieg, J.; Alkema, W., BioVenn – a web application for the comparison and visualization of biological lists using area-proportional Venn diagrams. BMC Genomics 2008, 9, (1), 488.

29. Gao, J.; Tarcea, V. G.; Karnovsky, A.; Mirel, B. R.; Weymouth, T. E.; Beecher, C. W.; Cavalcoli, J. D.; Athey, B. D.; Omenn, G. S.; Burant, C. F.; Jagadish, H. V., Metscape: a Cytoscape plug-in for visualizing and interpreting metabolomic data in the context of human metabolic networks. Bioinformatics 2010, 26, (7), 971–973.

30. Meier, F.; Brunner, A.-D.; Koch, S.; Koch, H.; Lubeck, M.; Krause, M.; Goedecke, N.; Decker, J.; Kosinski, T.; Park, M. A.; Bache, N.; Hoerning, O.; Cox, J.; Räther, O.; Mann, M., Online Parallel Accumulation–Serial Fragmentation (PASEF) with a Novel Trapped Ion Mobility Mass Spectrometer. Molecular & Cellular Proteomics 2018, 17, (12), 2534–2545.

31. Prianichnikov, N.; Koch, H.; Koch, S.; Lubeck, M.; Heilig, R.; Brehmer, S.; Fischer, R.; Cox, J., MaxQuant Software for Ion Mobility Enhanced Shotgun Proteomics*. Molecular & Cellular Proteomics 2020, 19, (6), 1058–1069.

32. Tyanova, S.; Cox, J., Perseus: A Bioinformatics Platform for Integrative Analysis of Proteomics Data in Cancer Research. In Cancer Systems Biology: Methods and Protocols, von Stechow, L., Ed. Springer New York: New York, NY, 2018; pp 133–148.

33. Huang, D. W.; Sherman, B. T.; Lempicki, R. A., Systematic and integrative analysis of large gene lists using DAVID bioinformatics resources. Nature Protocols 2009, 4, (1), 44–57.

34. Chong, J.; Soufan, O.; Li, C.; Caraus, I.; Li, S.; Bourque, G.; Wishart, D. S.; Xia, J., MetaboAnalyst 4.0: towards more transparent and integrative metabolomics analysis. Nucleic Acids Research 2018, 46, (W1), W486–W494.

35. Dettmer, K.; Nürnberger, N.; Kaspar, H.; Gruber, M. A.; Almstetter, M. F.; Oefner, P. J., Metabolite extraction from adherently growing mammalian cells for metabolomics studies: optimization of harvesting and extraction protocols. Analytical and Bioanalytical Chemistry 2011, 399, (3), 1127–1139.

36. Brown, K. A.; Chen, B.; Guardado-Alvarez, T. M.; Lin, Z.; Hwang, L.; Ayaz-Guner, S.; Jin, S.; Ge, Y., A photocleavable surfactant for top-down proteomics. Nature Methods 2019, 16, (5), 417–420.

37. Aballo, T. J.; Roberts, D. S.; Melby, J. A.; Buck, K. M.; Brown, K. A.; Ge, Y., Ultrafast and Reproducible Proteomics from Small Amounts of Heart Tissue Enabled by Azo and timsTOF Pro. Journal of Proteome Research 2021.

38. Meier, F.; Beck, S.; Grassl, N.; Lubeck, M.; Park, M. A.; Raether, O.; Mann, M., Parallel Accumulation–Serial Fragmentation (PASEF): Multiplying Sequencing Speed and Sensitivity by Synchronized Scans in a Trapped Ion Mobility Device. Journal of Proteome Research 2015, 14, (12), 5378–5387.

39. Meier, F.; Brunner, A.-D.; Koch, S.; Koch, H.; Lubeck, M.; Krause, M.; Goedecke, N.; Decker, J.; Kosinski, T.; Park, M. A.; Bache, N.; Hoerning, O.; Cox, J.; Räther, O.; Mann, M., Online Parallel Accumulation–Serial Fragmentation (PASEF) with a Novel Trapped Ion Mobility Mass Spectrometer *^&#x9;&#x9;^. Molecular & Cellular Proteomics 2018, 17, (12), 2534–2545.

40. Lopaschuk, G. D.; Jaswal, J. S., Energy Metabolic Phenotype of the Cardiomyocyte During Development, Differentiation, and Postnatal Maturation. Journal of Cardiovascular Pharmacology 2010, 56, (2).

41. Kolwicz Stephen, C.; Purohit, S.; Tian, R., Cardiac Metabolism and its Interactions With Contraction, Growth, and Survival of Cardiomyocytes. Circulation Research 2013, 113, (5), 603–616.

42. Reibel, D. K.; O’Rourke, B.; Foster, K. A.; Hutchinson, H.; Uboh, C. E.; Kent, R. L., Altered phospholipid metabolism in pressure-overload hypertrophied hearts. American Journal of Physiology-Heart and Circulatory Physiology 1986, 250, (1), H1–H6.

43. Forough, R.; Scarcello, C.; Perkins, M., Cardiac biomarkers: a focus on cardiac regeneration. The journal of Tehran Heart Center 2011, 6, (4), 179–186.

44. Scuderi, G. J.; Butcher, J., Naturally Engineered Maturation of Cardiomyocytes. Frontiers in Cell and Developmental Biology 2017, 5, 50.

45. Naba, A.; Clauser, K. R.; Ding, H.; Whittaker, C. A.; Carr, S. A.; Hynes, R. O., The extracellular matrix: Tools and insights for the “omics” era. Matrix Biology 2016, 49, 10–24.

46. Chang, C. W.; Dalgliesh, A. J.; López, J. E.; Griffiths, L. G., Cardiac extracellular matrix proteomics: Challenges, techniques, and clinical implications. PROTEOMICS – Clinical Applications 2016, 10, (1), 39–50.

47. Knott, S. J.; Brown, K. A.; Josyer, H.; Carr, A.; Inman, D.; Jin, S.; Friedl, A.; Ponik, S. M.; Ge, Y., Photocleavable Surfactant-Enabled Extracellular Matrix Proteomics. Analytical Chemistry 2020, 92, (24), 15693–15698.

