## Supplemental Information for "A Multi-omics Method Enabled by Sequential Metabolomics and Proteomics for Human Pluripotent Stem Cell-derived Cardiomyocytes" (from another version of this preprint: version 1, 2021-06-23)

### SUPPLEMENTAL INFORMATION: TABLE OF CONTENTS


***Section*** ***Pages***

Materials and Methods……………………………………………………………...…....….3-9

Supplemental Figures....………………………………………………………...…....……..10-24

Supplemental Tables....………………………………………………………...…....………25-28

Supplemental References……………………………………………………………...29

<u>**Materials and Methods**</u>

**Materials**

*Reagents and chemicals*: Unless otherwise noted, reagents and chemicals were purchased from Sigma-Aldrich, Inc. (St. Louis, MO, USA). HPLC-grade water, ethanol, and acetonitrile were purchased from Fischer Scientific (Fair Lawn, NJ, USA).

**hPSC and hPSC-CM Cell Culture**

Human induced pluripotent stem cells (WTC11 cell line) were maintained as described previously [1]. Briefly, cells were cultured in mTeSR1 (STEMCELL Technologies - 85850) on Growth Factor Reduced Matrigel (Corning – 354230)-coated 6-well plates (Corning COSTAR – 07-200-82) in a cell culture incubator (Sanyo MCO-18AC; 37°C, 5% CO2, 95% RH) and passaged with Versene (Life Technologies 15040066) at ~70% confluency (every 3-5 days).

hPSCs were differentiated into cardiomyocytes using the GiWi protocol as described previously [1]. Briefly, hPSCs were singularized in Accutase (Innovative Cell Technology - AT104), resuspended in mTeSR1 with 5 uM Y-27632 (Tocris - 1254), seeded onto Matrigel-coated 12-well plates, and incubated at room temperature for 30 min prior to transfer to a cell culture incubator on day minus (D-2) of differentiation; media was replaced with fresh mTeSR1 the following day (D-1). The subsequent day (D0), media was replaced with RPMI1640 (Life Technologies - 11875119) + 2% v/v B27 minus insulin (B27-; Life Technologies - 0050129SA) with the addition of 8 µM CHIR99021 (Selleckchem - S1263); 48 hours later (D2) media was replaced with B27- with the addition of 5 µM IWP2 (Tocris - 3533). On D4, media was replaced with fresh B27-; on D6, D8, D10, and every 3 days thereafter media was replaced with RPMI1640 + B27 supplement (Life Technologies - 17504044) until collection on day 42.

**Validation of hPSC-CM differentiation**

hPSC-CM differentiation efficiency was assessed in parallel wells by flow cytometry for cardiac troponin T (cTnT)-positive CMs on Day 16 of differentiation as described previously [1]. Briefly, cells were singularized in Accutase, fixed in 1% paraformaldehyde, and stored in 90% methanol at -20 °C until analysis. Approximately 200,000 cells were rinsed in flow buffer (FB = 0.5% w/v BSA in DPBS), incubated overnight at 4 °C in msIgG1-anti-cTnT primary antibody (ThermoFisher – MA5-12960) diluted 1:250 in FB + 0.1% v/v Triton X-100, rinsed with FB, incubated for 1 h at room temperature in AF488-anti-msIgG1 secondary antibody (ThermoFisher – A-21121) diluted 1:1000 in FB + 0.1% v/v Triton X-100, rinsed and resuspended in 300 µL FB prior to analysis on a BD Accuri C6 Plus flow cytometer. Undifferentiated hPSC negative control samples were stained in parallel. The resulting purity was 72 ± 2% cTnT+ (n=5); representative flow cytometry gating presented in **Figure S1**.

**Sequential Extraction of hPSC-CM Culture**

Metabolites and proteins were sequentially extracted using a one-step solvent-based quench technique [2]. Cell culture media was aspirated. Each hPSC-CM monolayer was washed twice with 2.0 mL $Ca^{2+}$ $Mg^{2+}$-free DPBS. Cells were incubated in 1.0 mL cold (4 ºC) 100% methanol for 1 minute to quench all cellular metabolic activity. Subsequently, cells were scraped into a 1.5 mL microcentrifuge tubes in methanol, which is referred to as “Metabolite-Protein Mixture.” Metabolite-Protein Mixture was vortexed for 10 seconds and incubated on a nutating mixer for 10 min at 4 ºC. The methanol-cell mixture was centrifuged for 5 min at 14,000 g, creating a metabolite-rich supernatant and protein-rich cell pellet. The pellet was rinsed with 200 µL DPBS. Metabolite and protein samples were dried under vacuum and stored at -80 ºC until further analysis.

**Untargeted Metabolomics of hPSC-CM**

*Metabolomics Sample Preparation:* Dried metabolite samples were reconstituted in 1:1 methanol:water prior to MS analysis by flow injection to a Bruker 12T solariX FTICR MS. Samples were thawed on ice for 1 min prior to the addition of cold methanol (100 µL). The sample was vortexed on high for 1-2 min. MS-grade water (100 µL) was added to the sample and vortexed on high for 1-2 min. An aliquot of 15 µL was taken from each sample to create a pooled Quality Control sample.

*Metabolomics Data Acquisition:* FIE-FTICR MS-based metabolomics fingerprinting was performed using a Waters ACQUITY UPLC M-Class System (Waters Corporation, Milford, MS, USA) coupled to a Bruker solariX 12T FTICR mass spectrometer (Bruker Daltonics, Bremen, Germany) without an LC column using a setup as described previously [3]. Electrospray ionization was used to introduce the samples to the mass spectrometer. MS-grade solvents were used to prepare mobile phases for positive (0.1% FA in 1:1 water/methanol) and negative (10 mM ammonium acetate in 1:1 water/methanol) ionization modes. A 5-minute isocratic hold of 100% pre-mixed mobile phase using a flow rate of 15 µL/min delivered the flow injection to the FTICR MS. A sample injection volume of 5.0 µL was used. Data acquisition parameters were used as described previously [4]. Briefly, 50 scans per spectrum were collected using an ion accumulation period of 0.1s per spectrum, totaling 3.5 min per sample run. The estimated resolving power at 400 *m/z* was 190,000. The FTICR MS was calibrated prior to data acquisition in each ionization mode using 5 mM NaTFA.

*Metabolomics Data Analysis:* MS data were processed using MetaboScape v 4.0 (Bruker Daltonics, Bremen, Germany). Bucket lists for data collected in positive and negative ionization modes were generated using the T-Rex 2D algorithm and combined to create one merged bucket list using a mass error tolerance of <1.0 ppm. A blank filtering step was applied, wherein features with an intensity of less than 3x of the intensity in the blank were filtered from the data set. SmartFormula annotations were performed in MetaboScape as described previously [4]. Features were annotated by accurate mass using the Mass Bank of North America and METLIN metabolite databases [5] with a maximum error tolerance of 3.0 ppm. Metabolite features were manually validated using DataAnalysis software (Bruker Daltonics, Bremen, Germany) and confirm MetaboScape feature detection. Overlap in metabolite features between hPSC-CMs was visualized using BioVenn [6]. Metabolites were mapped to networks using MetScape plugin in CytoScape v 3.8.1 [7].

**Global Proteomics Analysis of hPSC-CM**

*Proteomics Sample Preparation:* Proteins from the dried cell pellet were solubilized in 10 μL buffer containing 0.25% w/v Azo, a MS-compatible surfactant shown to increase enzymatic digestion efficiency [3, 8]. Samples underwent sonication and freeze-thaw cycles to aid protein solubilization and were diluted to 0.05% w/v Azo before proceeding. Samples were normalized using Bradford assay, reduced using 10 mM DTT for 1 hour, alkylated using 20 mM IAA for 30 minutes, and digested in a 50:1 protein:enzyme ratio using Trypsin Gold (Promega, Madison,

Wisconsin, USA) for 3+ hours. After quenching enzymatic activity using formic acid, samples were irradiated at 305 nm for 5 min and desalted using C18 ZipTips. Peptides were dried under vacuum and reconstituted in 0.1% formic acid in water at 0.2 mg/mL. Peptide concentration was analyzed using a NanoDrop 2000 prior to LC-MS analysis.

*Global Proteomics Data Acquisition:* Nanoflow LC-MS analysis of tryptic peptides was performed using a timsTOF Pro (Bruker Daltonics, Bremen, Germany) [9] coupled to a nanoElute nano-flow UHPLC system (Bruker Daltonics, Bremen, Germany) using a Captivespray nano-electrospray ion source. Approximately 200 ng peptides were loaded onto an IonOpticks capillary C18 column (25 cm x 75 µm i.d., 1.7 µm) for each injection. Peptides were separated at 55 ºC using a 400 nL/min flow rate and a stepwise gradient of 2 to 17% B over 60 minutes and 17 to 25% B from 60 to 90 min, followed by a 30 min wash step at 85% B, totaling 120 min per run. Mobile phases A and B were 0.1% formic acid in water and 0.1% formic acid in acetonitrile, respectively. The timsTOF Pro was operated in Parallel Accumulation and Serial Fragmentation (PASEF) mode [9]. The mass range for MS and MS/MS scans was 100-1700 *m/z*. Ion mobility resolution was set to 0.60-1.60 V*s/cm over a ramp time of 100 ms. Data-dependent acquisition was performed using 10 PASEF MS/MS scans per cycle using a locked duty cycle of 100%. An active exclusion time of 0.4 min was applied to precursors that reached 20,000 intensity units and reconsider them after 4.0 min. Collisional induced dissociation (CID) energies were scaled on ion mobilities, 1/k0. Peptide loading was assessed by comparing the intensity of the total ion current to 200 ng of a human K562 whole cell lysate standard (Promega, Madison, Wisconsin, USA) prepared at 0.2 µg/µL.

*Proteomics Data Analysis:* Tandem mass spectra were searched against the UniProt human database UP000005640 (accessed 04 September 2020) using MaxQuant version 1.6.17.0 [10]. A

1% false discovery rate was used at the peptide and protein level. TIMS-DDA runs were set half-width to 6 and resolution to 32,000. The maximum mass tolerance for precursor ions was 30 ppm. Peptides had a minimum of 7 amino acids with carbamidomethylation as a fixed modification and N-terminal acetylation and methionine oxidation as variable modifications. Enzyme specificity was set to trypsin/P and a maximum of 2 missed cleavages were allowed. Match between runs feature was used with a 0.7 min window after retention time alignment to maximize identifications between runs. Label free quantification (LFQ) was performed using classic normalization and a minimum ratio count of 2. The match between runs feature was used with a maximum retention window of 0.7 minutes and the ion mobility window set to 0.05 V*s/cm. MS/MS spectra were required for LFQ comparisons.

Data was further processed using Perseus software v. 1.6.14.0 [11]. The dataset was annotated for Gene Ontology Cellular Compartment (GOCC), Gene Ontology Biological Process (GOBP), Gene Ontology Molecular Function (GOMF), and the Kyoto Encyclopedia for Genes and Genomes (KEGG) for each identified protein group. Potential contaminants and reverse hits were excluded. To determine correlation of LFQ intensities across samples, Perseus software was used to transform the data by log2x and generate scatter plots with Pearson correlation coefficients. Overlap in protein groups and peptides identified between hPSC-CMs was visualized using BioVenn [6]. Pathway analysis and functional enrichment was performed on the dataset using the Database for Annotation, Visualization, and Integrated Discovery (DAVID) [12].

**Multi-omics (Metabolomics-Proteomics) Data Integration**

To enable pathway analysis, KEGG identifiers were assigned to metabolites annotated from METLIN and MoNA databases. Protein (UniProt identifications) and metabolite (KEGG identifiers) lists were uploaded to MetaboAnalyst 5.0 for integrative pathway analysis [13]. The Joint Pathway Analysis module was used to determine significantly enriched pathways within the dataset (hypergeometric test, $p \leq 0.05$). Pathways considered abundantly represented in the dataset were selected for granular visualization of protein and metabolite populations using MetaboAnalyst network visualization tools. The Network Explorer module was used to visualize enriched metabolic-related pathways overlaid on the KEGG metabolic pathway map (ko01100).

**Supplemental Figures**

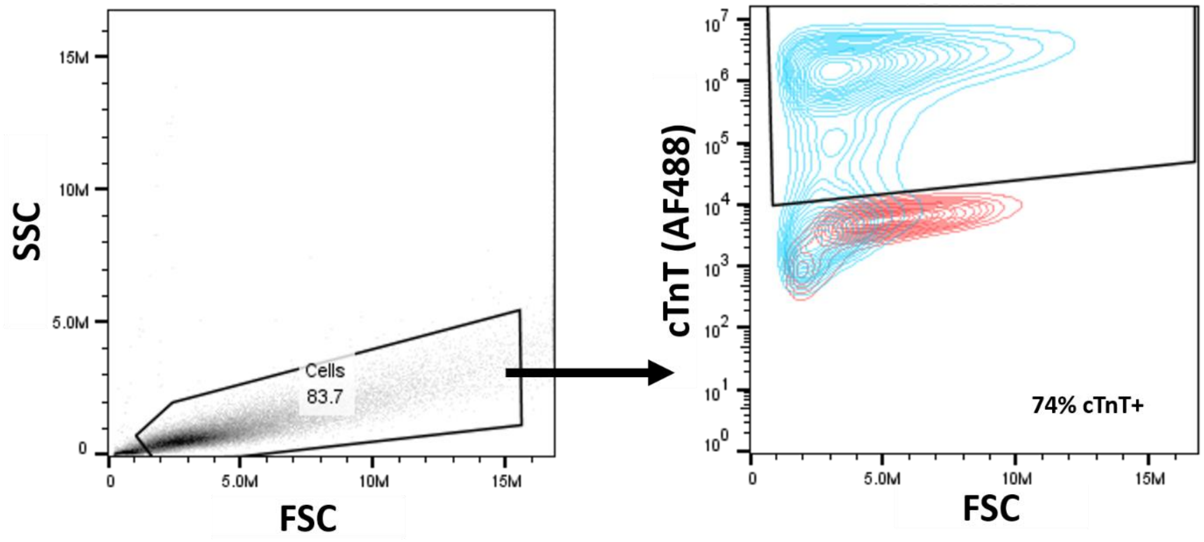


**Figure S1. hPSC-CM purity validation.** Flow cytometry gating of Day 16 samples assessed for presence of cardiac troponin T; gating of cells on FSC/SSC axes (left). Resulting cells then gated on FSC/FL-1 for % CMs based on cTnT+ staining compared to stem cell negative control sample (red).

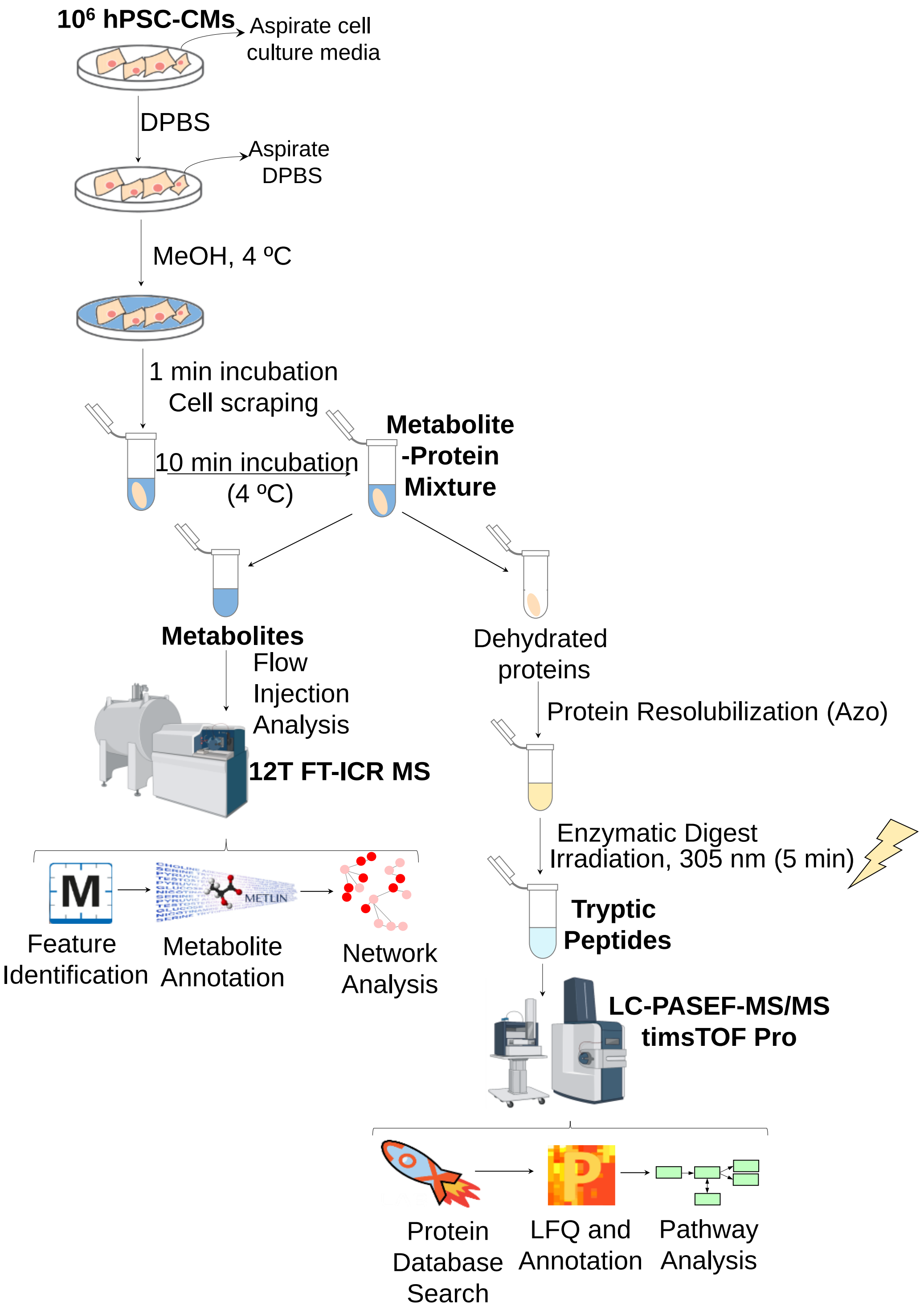


**Figure S2.** Schematic of sequential extraction of both metabolite and proteins from hPSC-CM for integrated metabolomics and proteomics. Detailed extraction procedure and analysis is described in **Supplemental Materials and Methods**.

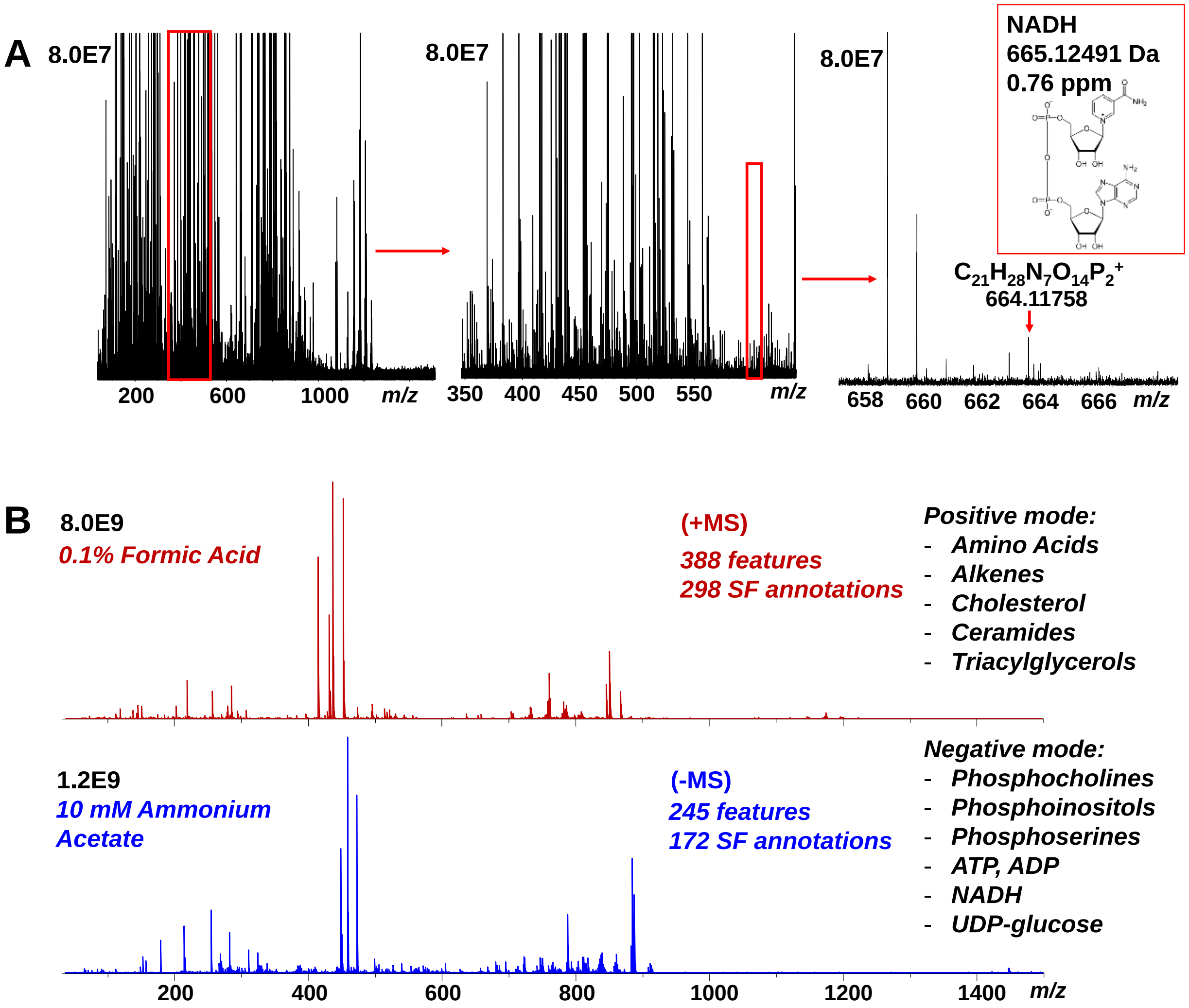


**Figure S3. High Resolution and Rapid Polarity Switching of the FIE-FTICR MS Platform Enables Identification of Hundreds of Metabolites per Sample. A)** Broadband high resolution metabolite performance can resolve thousands of metabolite features from complex spectra with high resolution and mass accuracy, enabling the identification of hundreds of metabolites with high mass accuracy, including NADH with sub-ppm mass error (0.76 ppm). **B)** Integrated FIE-FTICR MS platform for both metabolomics and lipidomics. Rapid polarity switching on the FIE-FTICR MS platform enables the detection of unique and nonredundant metabolites from each polarity mode, including classical small molecule metabolites and many classes of phospholipids.

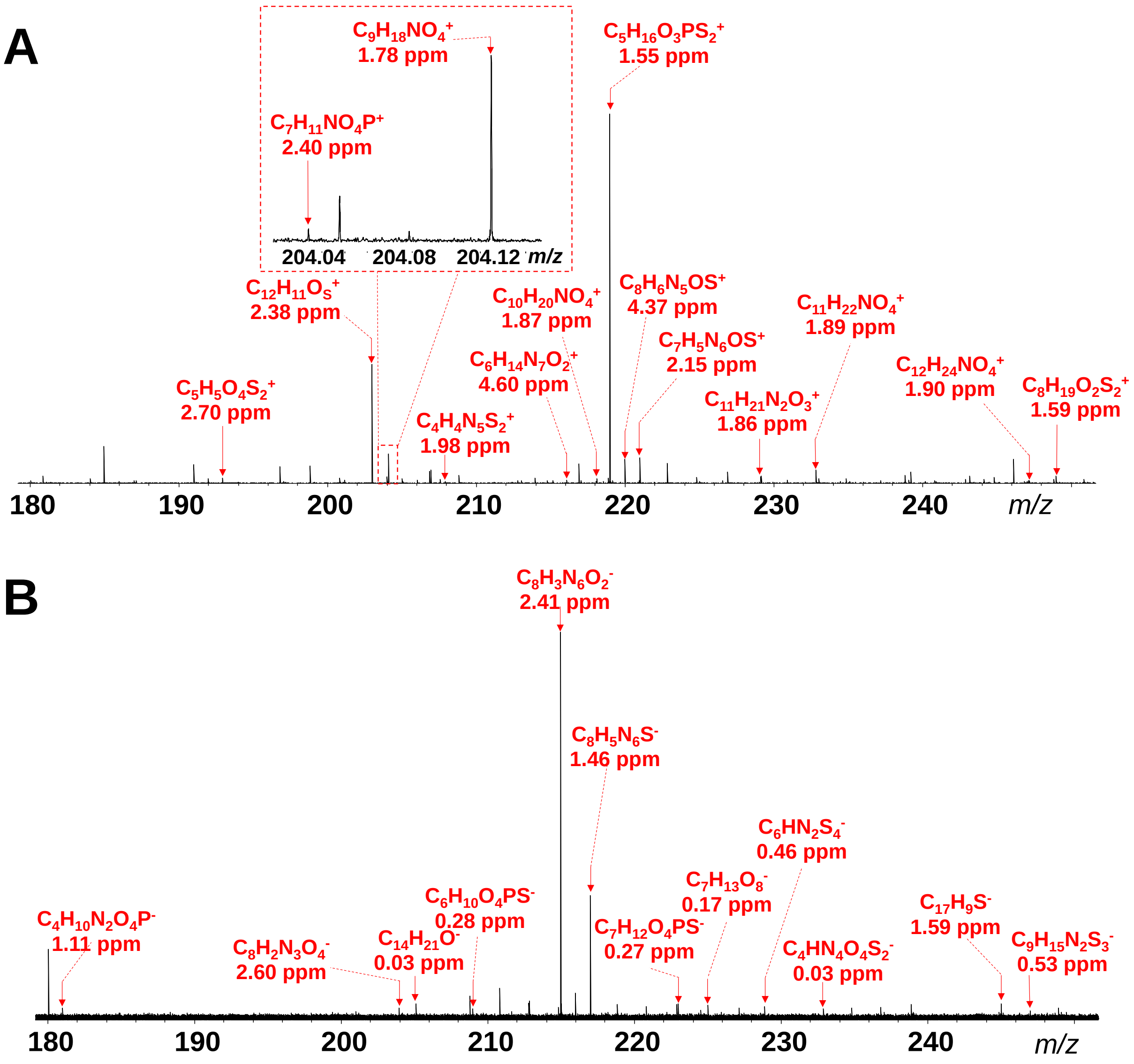


**Figure S4. Representative broadband metabolite spectra show ultrahigh mass accuracy of SmartFormula annotations in positive and negative ionization modes. A)** A representative zoomed in positive ion mode mass spectrum reveals 14 unique features in a 70 m/z range annotated within 5 ppm error using the SmartFormula (SF) function, with 12 of them annotated around 2 ppm. **B)** A representative zoomed in negative ion mode mass spectrum revealed 12 unique features in a 70 m/z range annotated within 5 ppm error using the SF function, with 7 of them annotated under 1 ppm.

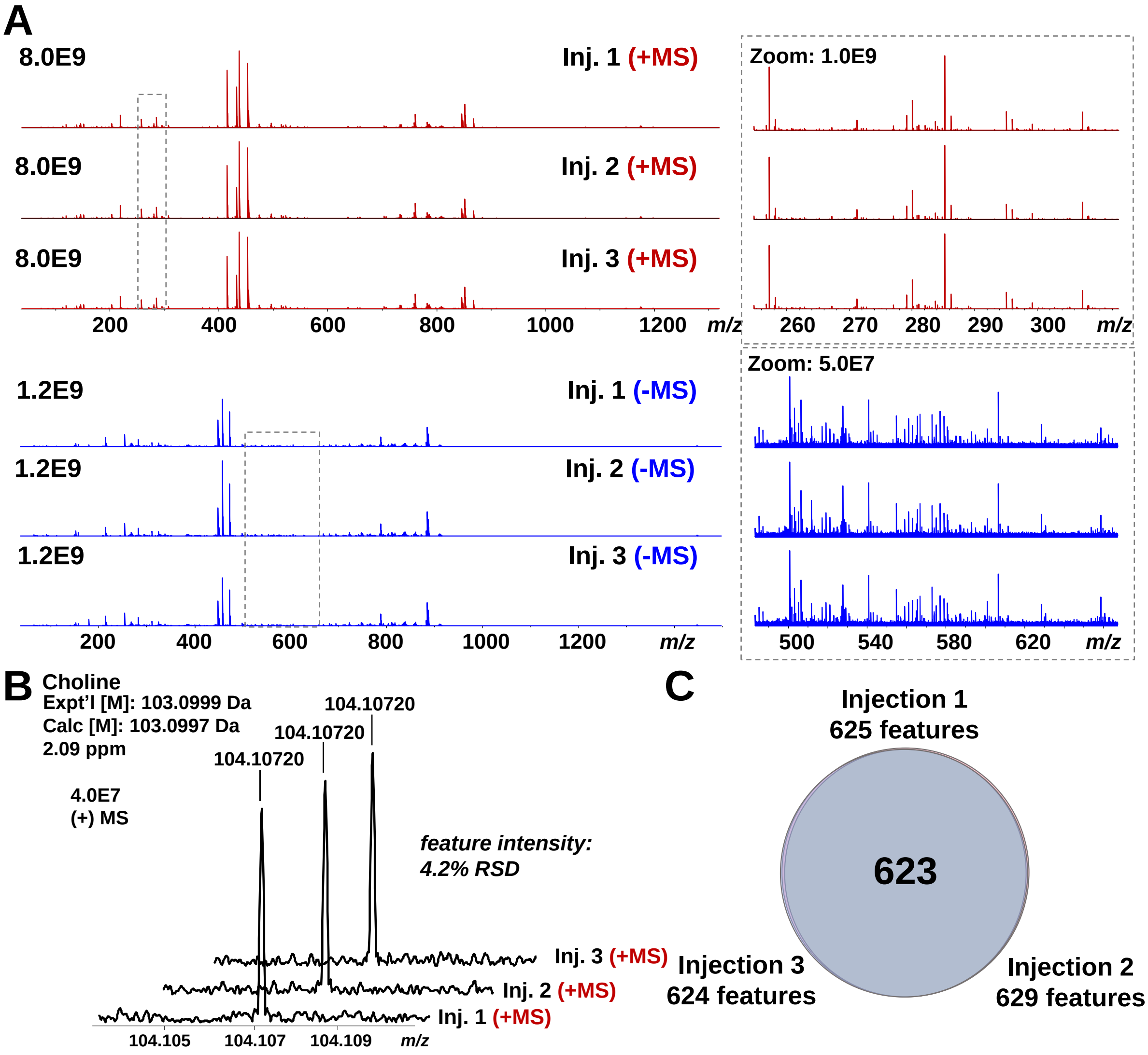


**Figure S5. FIE-FTICR MS Platform Enables Rapid and Reproducible Collection of Complex Metabolite Spectra**. **A)** Highly reproducible experiments shown by similarities in mass spectra in metabolite feature intensities were obtained for both positive (top, red) and negative (blue, below) ionization modes. A zoomed panel in each ionization mode shows highly similar metabolite spectra for lower-abundance features. **B)** Choline, a central ATP-generating metabolite, is identified between triplicate injections with and 4.2% relative standard deviation (RSD) between triplicate injections. **C)** Injection replicates showed 99% overlap in features detected, with 623 features shared between triplicate injections and 630 total.

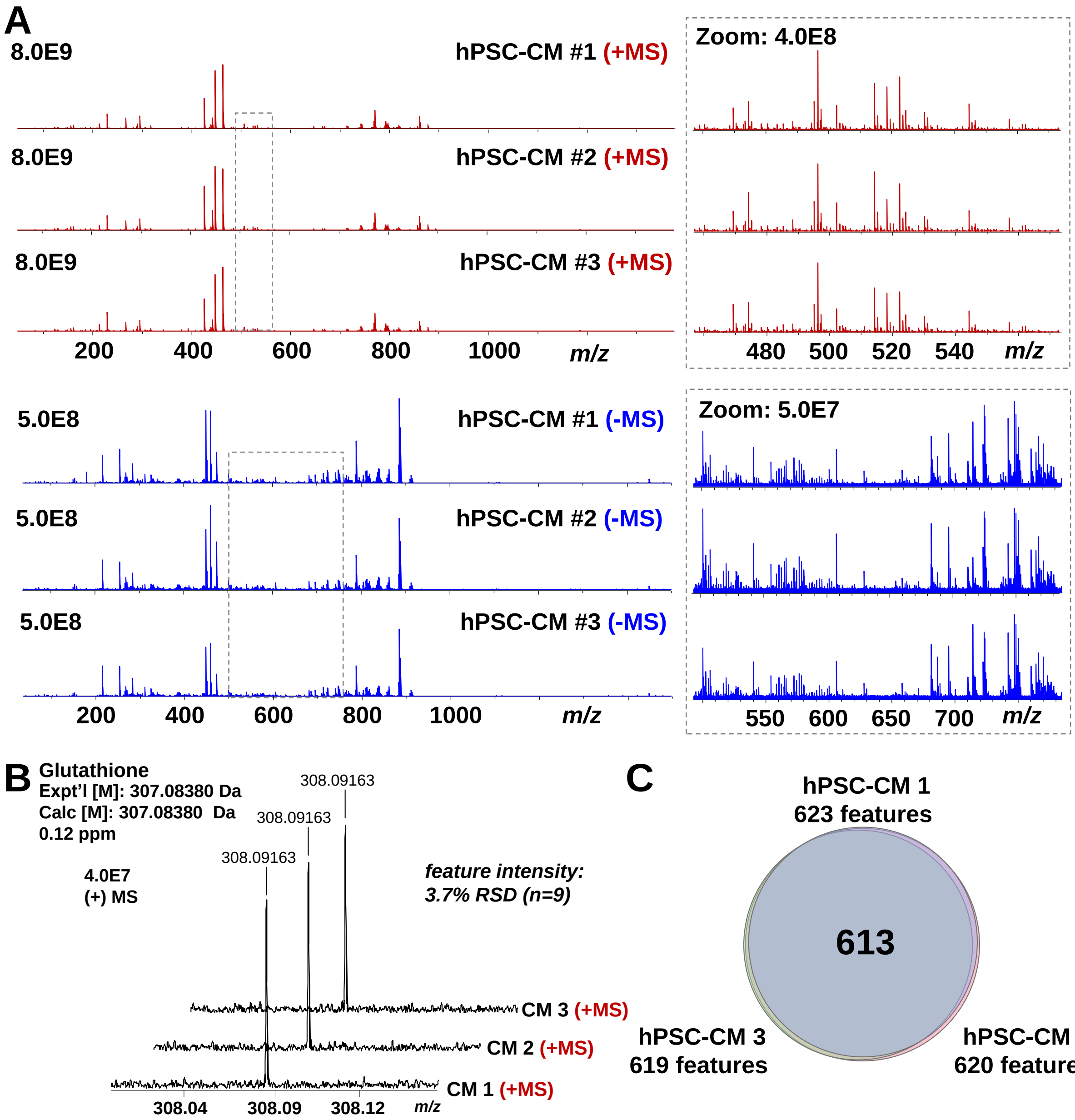


**Figure S6. hPSC-CM Extraction Replicates Yield Highly Reproducible Mass Spectra.** FIE-FTICR MS metabolite mass spectra for n=3 hPSC-CM extraction replicates are highly reproducible in in **A)** positive ionization mode and negative ionization mode, with zoom inlays for each to show lower-abundant features in detail. **B)** Glutathione, a coenzyme involved in amino acid metabolism, is identified between hPSC-CM biological replicates injections with no deviation in *m/z* and only 3.7% RSD in intensity between triplicate injections for each CM (n=9 measurements). **C)** Overlap of features detected between hPSC-CM replicates. Features identified in 3 of 3 injections were included for each biological replicate. 613 features were shared between three hPSC-CM biological replicates.

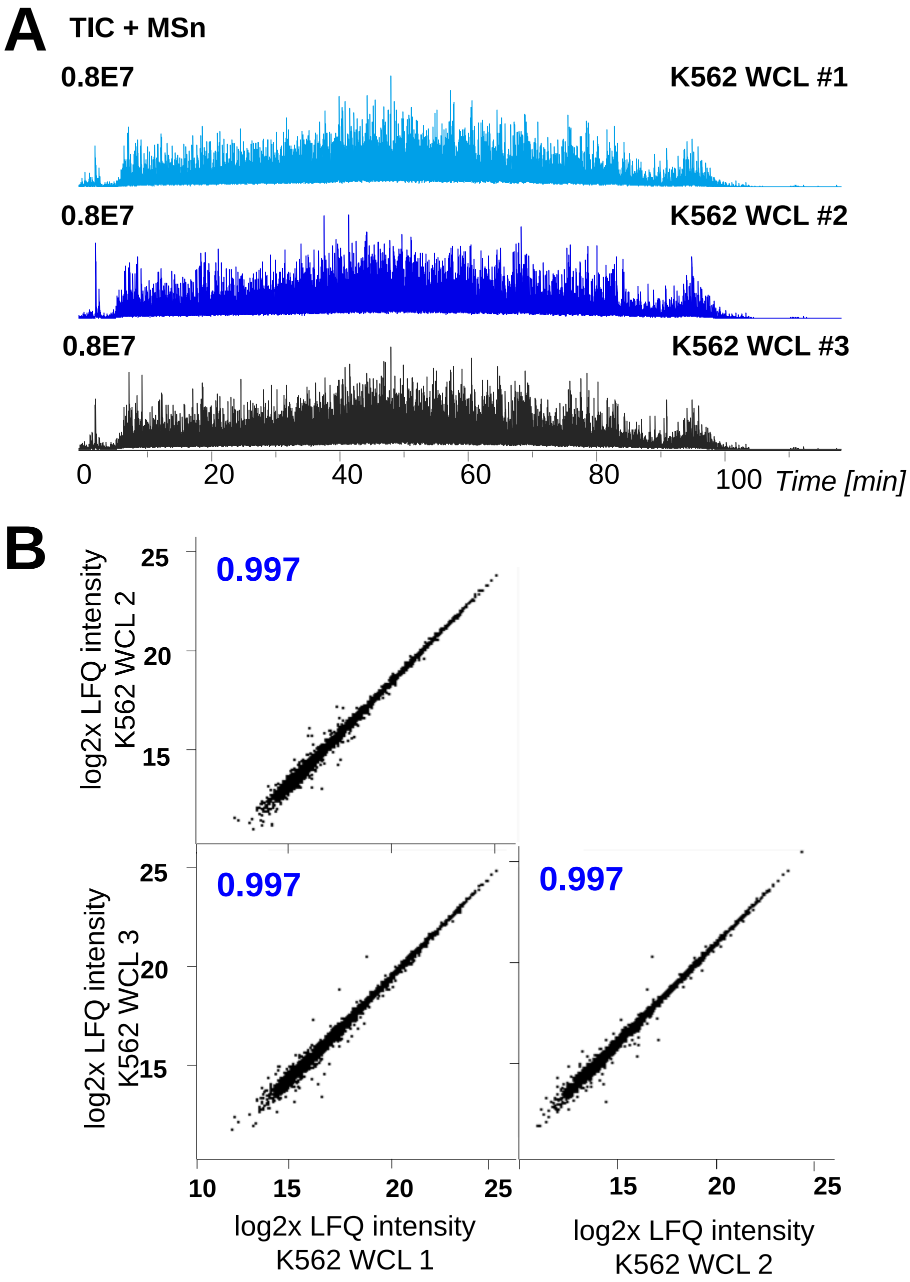


**Figure S7. K562 Whole Cell Lysate (WCL) bracketing standard injections show instrument robustness across 58 h sample run. A)** Reproducible TIC+MSn spectra for each WCL standard injection show similar profile and intensity throughout the sample run. **B)** High correlation of LFQ intensities for each standard injection is observed, with the Pearson correlation coefficient = 0.997 for each comparison.

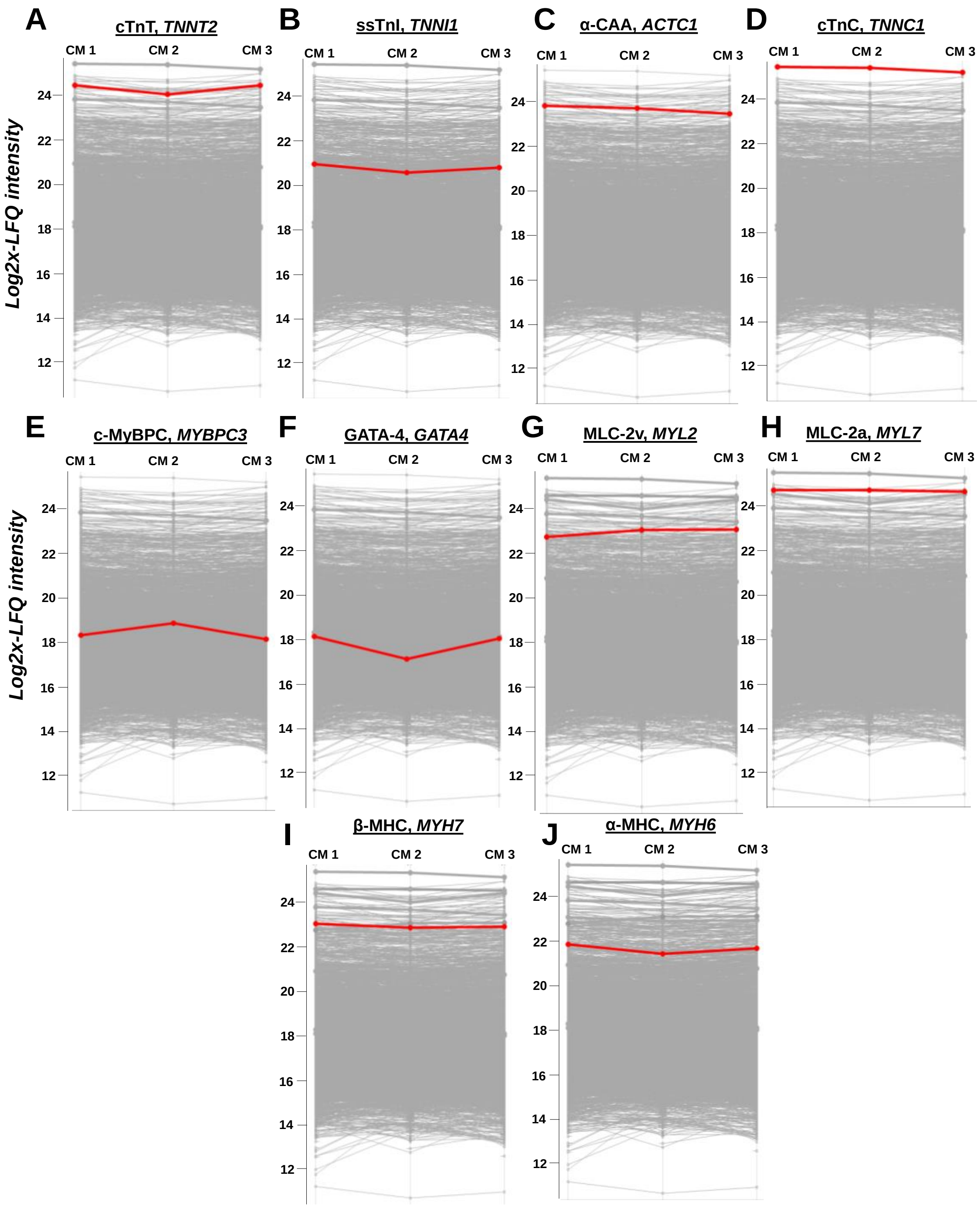


**Figure S11. Profile plots of cardiac proteins found in hPSC-CM biological replicates (CMs 1-3) including A)** cardiac troponin T, cTnT (gene: *TNNT2*) **B)** slow skeletal troponin I, ssTnI (gene: *TNNI1*), **C)** α-cardiac muscle actin, α-CAA (gene: *ACTC1*), **D)** cardiac troponin C, cTnC (gene: *TNNC1*), **E)** cardiac myosin binding protein C, c-MyBPC (gene: *MYBPC3*), **F)** GATA binding protein 4, GATA-4 (gene: *GATA4*), **G)** myosin light chain, ventricular isoform, MLC-2v (gene: MYL2), H) myosin light chain, atrial isoform, MLC-2a (gene: MYL7), **I)** beta-myosin heavy chain, β-MHC (gene: MYH7), and **J)** alpha-myosin heavy chain, α-MHC (gene: MYH6). Results generated by Perseus software; log2x-transformed LFQ intensities shown.

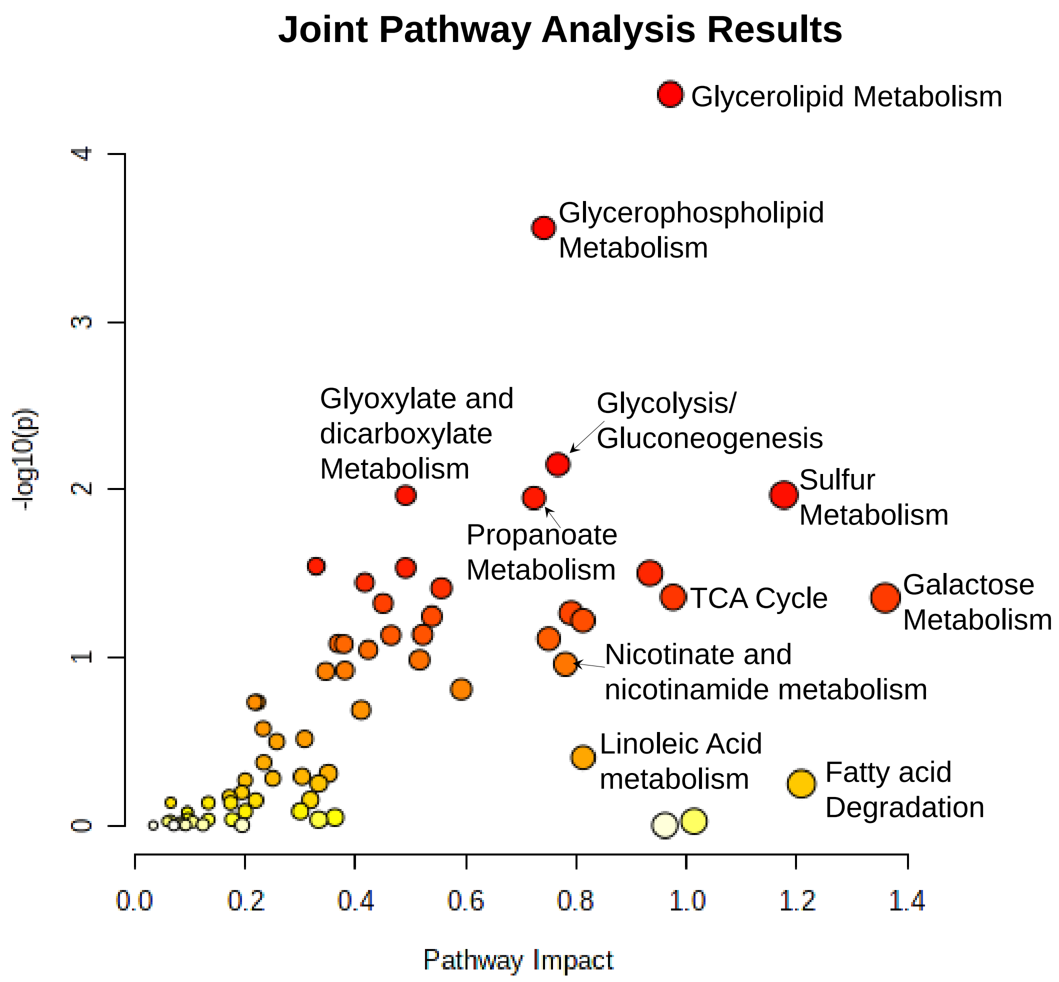


**Figure S12. Joint Pathway Analysis Results from MetaboAnalyst**. Metabolic pathways abundantly represented in the dataset (y-axis, determined by hypergeometric test where $p \leq 0.05$) are measured by pathway impact (x-axis, corresponding to the representation of proteins/metabolites identified in each pathway). Some metabolic pathways considered significantly enriched were glycerolipid metabolism, glycerophospholipid metabolism, glycolysis or gluconeogenesis, Sulfur metabolism, Glyoxylate and dicarboxylate metabolism, Propanoate metabolism, Aminoacyl-tRNA biosynthesis, and Glutathione metabolism. See **Table S5** for complete list of significantly enriched pathways. Colors from yellow to red indicate different levels of significance.

**Supplemental Tables**

**Table S1**. Representative metabolites identified in hPSC-CMs. Seven metabolite categories, including acyl-carnitines, amino acids and derivatives, triglycerides, glycerophospholipids, and sphingolipids are highlighted with representative metabolites listed. For full list, refer to **Supplemental File 2.**

| Identified | Calc Mass (Da) | Expt'l Mass (Da) | m/z | Mass error (ppm) | PubChem ID |
|---|---|---|---|---|---|
| **Metabolites** | | | | | |
| Adenosine triphosphate (ATP) | 506.9957 | 506.9958 | 505.9885 | 0.0 | 5957 |
| Adenosine diphosphate (ADP) | 427.0294 | 427.0294 | 426.0221 | 0.0 | 6022 |
| Choline | 103.0997 | 103.0999 | 104.1072 | 2.1 | 305 |
| Glutathione | 307.0838 | 307.0838 | 306.0766 | 0.1 | 745 |
| NAD+ | 663.1091 | 663.1103 | 664.1176 | 1.8 | 93556 |
| NADH | 665.1248 | 665.1249 | 664.1176 | 0.2 | 5893 |
| Oleandomycin 2'-O-phosphate | 767.3857 | 767.3843 | 766.3770 | 1.8 | 440123 |
| Glucose | 180.0634 | 180.0638 | 203.0530 | 2.2 | 5793 |
| **Fatty Acids** | | | | | |
| Arachidonaic acid | 304.2402 | 304.2405 | 102.4208 | 0.9 | 444899 |
| Eicosatrienoic acid | 306.2556 | 306.2548 | 307.2621 | 2.4 | 5280581 |
| DGDG 32:2 | 888.5810 | 888.5834 | 887.5762 | 2.7 | - |
| DGTS 36:4 | 759.6013 | 759.6014 | 760.6086 | 0.1 | 363612530 |
| Diethylhexyl adipate | 370.3083 | 370.3090 | 393.2982 | 1.8 | 7641 |
| L-Carnitine | 161.1052 | 161.1055 | 162.1128 | 1.9 | 10917 |
| Linoleic acid | 280.2402 | 280.2402 | 279.2330 | 0.0 | 5282457 |
| **Acyl Carnitines** | | | | | |
| Propionyl-L-carnitine | 217.1314 | 217.1318 | 218.1391 | 1.9 | 107738 |
| Oleoyl-L-Carnitine | 425.3505 | 425.3513 | 464.3144 | 1.8 | 46907933 |
| Isovalerylcarnitine | 245.1627 | 245.1632 | 246.1704 | 1.8 | 6426851 |
| L-Acetylcarnitine | 203.1158 | 203.1161 | 204.1234 | 1.9 | 7045767 |
| L-Hexanoylcarnitine | 259.1784 | 259.1789 | 260.1862 | 2.0 | 3246938 |
| Sphingolipids | | | | | |
| HexCer-AP t34:2 | 729.5391 | 729.5396 | 730.5469 | 0.7 | 384018563 |
| SHexCer d35:0 | 795.5530 | 795.5550 | 794.5477 | 2.5 | 383960910 |
| SHexCer d35:1 | 793.5413 | 793.5395 | 792.5322 | 2.3 | 383890186 |
| SM d32:1 | 674.5363 | 674.5375 | 675.5448 | 1.8 | 11433862 |
| SM d34:1 | 702.5676 | 702.5688 | 703.5760 | 1.7 | 9939941 |
| SM d34:2 | 700.5519 | 700.5531 | 701.5604 | 1.7 | 154573092 |
| **Amino Acids and derivatives** | | | | | |
| 5-Oxoprolinate | 129.0426 | 129.0429 | 152.0321 | 2.3 | 499 |

| | | | | | |
|---|---|---|---|---|---|
| L-Arginine | 174.1117 | 174.1120 | 175.1193 | 2.0 | 6322 |
| Leucylproline | 228.1474 | 228.1478 | 229.1551 | 1.8 | 80817 |
| L-Glutamine | 146.0691 | 146.0694 | 169.0587 | 2.1 | 5961 |
| D-Alanine | 89.0477 | 89.0480 | 128.0111 | 3.5 | 239 |
| **Glycerophospholipids** | | | | | |
| Glycerophosphocholine | 257.1028 | 257.1028 | 280.0926 | 0.1 | 657272 |
| LPC 16:0 | 495.3325 | 495.3318 | 496.3407 | 1.5 | 460602 |
| LPS 21:0 | 567.3536 | 567.3536 | 566.3464 | 0.0 | 75959202 |
| PEtOH 50:3 | 922.7391 | 922.7411 | 923.7484 | 2.2 | 383914791 |
| PG 36:2 | 774.5410 | 774.5414 | 773.5341 | 0.5 | 52927248 |
| PG 40:7 | 820.5254 | 820.5260 | 819.5187 | 0.7 | 53480628 |
| PC 31:2e | 701.5359 | 701.5362 | 700.5289 | 0.3 | 5283541 |
| PC 33:1 | 745.5622 | 745.5625 | 744.5552 | 0.4 | 52922442 |
| PC 35:4e | 753.5672 | 753.5675 | 752.5602 | 0.3 | 24779288 |
| PE 34:1 | 717.5309 | 717.5311 | 716.5238 | 0.4 | 5283496 |
| PE 36:4e | 725.5359 | 725.5362 | 724.5289 | 0.3 | 42607456 |
| PE 36:5e | 723.5203 | 723.5205 | 722.5133 | 0.3 | 52924155 |
| PI 34:1 | 836.5415 | 836.5414 | 835.5342 | 0.1 | 71296232 |
| PI 34:2 | 834.5258 | 834.5263 | 833.5191 | 0.6 | 11072889 |
| PI 36:1 | 864.5728 | 864.5732 | 863.5659 | 0.5 | 52927527 |
| PS 34:1 | 761.5206 | 761.5209 | 760.5137 | 0.4 | 52926066 |
| PS 35:1 | 775.5363 | 775.5365 | 774.5292 | 0.2 | 52925642 |
| PS 36:0 | 791.5676 | 791.5705 | 790.5608 | 3.7 | 9547096 |
| Dioleoyl phosphatidylethanolamine | 743.5465 | 743.5467 | 742.5395 | 0.3 | 6437392 |
| CL 70:7 | 1422.9566 | 1422.9571 | 710.4713 | 0.4 | 53480379 |
| CL 72:8 | 1448.9722 | 1448.9728 | 723.4791 | 0.4 | 5283539 |
| BMP 36:3 | 772.5254 | 772.5257 | 771.5185 | 0.4 | 383897955 |
| **Glycerolipids** | | | | | |
| TAG 54:5 | 880.7520 | 880.7527 | 881.7587 | 0.8 | 131756117 |
| TAG 54:7 | 876.7207 | 876.7202 | 877.7275 | 0.5 | 131756126 |
| TAG 56:6 | 906.7676 | 906.7683 | 907.7744 | 0.8 | 9544615 |

**Table S2. Diversity of Metabolites Identified in hPSC-CM Cell Culture.** Eleven metabolite categories highlighted from 205 total metabolites annotated from experimental databases with an error tolerance of ≤ 3.0 ppm.

| **Metabolite Group** | **# Metabolites** |
|---|---|
| Central Metabolites | 26 |
| Fatty Acids | 10 |
| Acyl Carnitines | 4 |
| Sphingolipids | 6 |
| Amino Acids | 8 |
| Glycerolipids | 9 |
| Cardiolipins | 3 |
| Phosphotidylserines | 14 |
| Phosphatidylglycerols | 3 |
| Phosphatidylcholines | 23 |
| Phosphatidylethanolamines | 27 |

**Table S3. Reproducible protein yields from hPSC-CM cell pellet after sequential extraction.** Total protein extracted per hPSC-CM well as determined by Bradford Assay (Coomassie Plus, Thermo Fischer Scientific). Assay was performed as directed in manual; Bovine Serum Albumin used as calibration standard; $R^2$ = 0.996 from 0-0.25 mg/mL. Additionally, we determined the presence of the anionic surfactant at low concentrations (0.001-0.0625% w/v) minimally affected the assay results (**Table S3**). The working concentration of the samples was 0.005% w/v due to a sample dilution factor of 10x prior to assay. SE: standard deviation.

| **Sample** | **µg/µL protein** | **µg protein total** |
|---|---|---|
| hPSC-CM #1 | 0.55 | 27 |
| hPSC-CM #2 | 0.52 | 26 |
| hPSC-CM #3 | 0.63 | 31 |
| *Average ± SE* | *0.57 ± 0.05* | *28 ± 3* |

**Table S4. Azo interferes with Coomassie Plus Bradford Assay (BFA) at 0.125% w/v.** A serial dilution of Azo from 0.5% to 0.001% in 25 mM ABC (aq) was used to test absorptive interference of Azo in BFA. Signal interference* begins between 0.0625% and 0.125% w/v Azo. We determined the anionic surfactant Azo had little to no effect on the Bradford Assay at our working concentration of 0.005% w/v due to a sample dilution factor of 10x prior to assay.

| % w/v Azo-6 | AU |
|---|---|
| 0.00 | 0.261* |
| 0.50 | 0.348* |
| 0.25 | 0.334* |
| 0.125 | 0.281* |
| 0.0625 | 0.259 |
| 0.0313 | 0.257 |
| 0.0156 | 0.256 |
| 0.00781 | 0.256 |
| 0.00391 | 0.255 |
| 0.00195 | 0.2595 |

**Table S5.** Top 25 most abundant KEGG pathways (as determined by FDR-corrected $p \leq 0.05$) identified using hPSC-CM proteomics data using the Database for Annotation, Visualization, and Integrated Discovery (DAVID). For full list, refer to Supplemental Table from DAVID (Supplemental File 4).

| KEGG Pathway Name | Gene Count | Gene Ratio (%) | FDR |
|---|---|---|---|
| hsa03040:Spliceosome | 111 | 2.62 | 5.76E-36 |
| hsa01130:Biosynthesis of antibiotics | 129 | 3.04 | 6.45E-20 |
| hsa03010:Ribosome | 91 | 2.14 | 9.06E-18 |
| hsa01200:Carbon metabolism | 80 | 1.89 | 9.06E-18 |
| hsa03013:RNA transport | 102 | 2.40 | 1.45E-14 |
| hsa04141:Protein processing in endoplasmic reticulum | 97 | 2.29 | 1.10E-12 |
| hsa04144:Endocytosis | 126 | 2.97 | 1.32E-12 |
| hsa00280:Valine, leucine and isoleucine degradation | 39 | 0.92 | 7.06E-12 |
| hsa05016:Huntington's disease | 103 | 2.43 | 3.63E-11 |
| hsa03050:Proteasome | 36 | 0.85 | 1.24E-10 |
| hsa05012:Parkinson's disease | 81 | 1.91 | 1.59E-10 |
| hsa05130:Pathogenic Escherichia coli infection | 39 | 0.92 | 4.43E-10 |
| hsa04510:Focal adhesion | 105 | 2.47 | 8.88E-10 |
| hsa01230:Biosynthesis of amino acids | 48 | 1.13 | 3.07E-09 |
| hsa00190:Oxidative phosphorylation | 74 | 1.74 | 5.05E-09 |
| hsa05010:Alzheimer's disease | 88 | 2.07 | 5.41E-09 |
| hsa03015:mRNA surveillance pathway | 55 | 1.30 | 2.12E-08 |
| hsa00020:Citrate cycle (TCA cycle) | 24 | 0.57 | 1.20E-06 |
| hsa05205:Proteoglycans in cancer | 88 | 2.07 | 9.84E-05 |
| hsa05100:Bacterial invasion of epithelial cells | 42 | 0.99 | 1.14E-04 |
| hsa00030:Pentose phosphate pathway | 21 | 0.49 | 1.14E-04 |
| hsa00630:Glyoxylate and dicarboxylate metabolism | 20 | 0.47 | 1.14E-04 |
| hsa00620:Pyruvate metabolism | 26 | 0.61 | 1.16E-04 |
| hsa01100:Metabolic pathways | 417 | 9.83 | 1.54E-04 |
| hsa00010:Glycolysis / Gluconeogenesis | 37 | 0.87 | 1.83E-04 |

**Table S6**. Top 20 significantly-enriched KEGG pathways from joint protein and metabolite list input, as determined by FDR-corrected $p \leq 0.05$. Pathway analysis performed by MetaboAnalyst 5.0. For full list, refer to Supplemental Table from MetaboAnalyst (Supplemental File 5).

| KEGG Pathway Name | FDR |
|---|---|
| Ribosome | 4.9E-32 |
| Spliceosome | 1.4E-20 |
| Oxidative phosphorylation | 2.2E-15 |
| Parkinson disease | 8.5E-14 |
| Huntington disease | 1.3E-12 |
| Thermogenesis | 1.3E-12 |
| RNA transport | 3.6E-12 |
| Alzheimer disease | 5.0E-10 |
| Non-alcoholic fatty liver disease (NAFLD) | 5.6E-07 |
| RNA degradation | 6.6E-06 |
| Phagosome | 7.7E-06 |
| Protein processing in endoplasmic reticulum | 1.1E-05 |
| Proteasome | 1.2E-05 |
| Cardiac muscle contraction | 1.2E-05 |
| Pathogenic Escherichia coli infection | 2.2E-05 |
| Lysosome | 4.0E-05 |
| Endocytosis | 2.6E-04 |
| Focal adhesion | 2.6E-04 |
| mRNA surveillance pathway | 5.9E-04 |
| Hypertrophic cardiomyopathy (HCM) | 7.7E-04 |

### Supplemental References


[1] X. Lian, J. Zhang, S.M. Azarin, K. Zhu, L.B. Hazeltine, X. Bao, C. Hsiao, T.J. Kamp, S.P. Palecek, Directed cardiomyocyte differentiation from human pluripotent stem cells by modulating Wnt/β-catenin signaling under fully defined conditions, Nature Protocols 8(1) (2013) 162-175.

[2] Q. Teng, W. Huang, T.W. Collette, D.R. Ekman, C. Tan, A direct cell quenching method for cell-culture based metabolomics, Metabolomics 5(2) (2008) 199.

[3] K.A. Brown, T. Tucholski, C. Eken, S. Knott, Y. Zhu, S. Jin, Y. Ge, High-Throughput Proteomics Enabled by a Photocleavable Surfactant, Angewandte Chemie International Edition 59(22) (2020) 8406-8410.

[4] Y. Zhu, B. Wancewicz, M. Schaid, T.N. Tiambeng, K. Wenger, Y. Jin, H. Heyman, C.J. Thompson, A. Barsch, E.D. Cox, D.B. Davis, A.R. Brasier, M.E. Kimple, Y. Ge, Ultrahigh-Resolution Mass Spectrometry-Based Platform for Plasma Metabolomics Applied to Type 2 Diabetes Research, Journal of Proteome Research 20(1) (2021) 463-473.

[5] C. Guijas, J.R. Montenegro-Burke, X. Domingo-Almenara, A. Palermo, B. Warth, G. Hermann, G. Koellensperger, T. Huan, W. Uritboonthai, A.E. Aisporna, D.W. Wolan, M.E. Spilker, H.P. Benton, G. Siuzdak, METLIN: A Technology Platform for Identifying Knowns and Unknowns, Analytical Chemistry 90(5) (2018) 3156-3164.

[6] T. Hulsen, J. de Vlieg, W. Alkema, BioVenn – a web application for the comparison and visualization of biological lists using area-proportional Venn diagrams, BMC Genomics 9(1) (2008) 488.

[7] J. Gao, V.G. Tarcea, A. Karnovsky, B.R. Mirel, T.E. Weymouth, C.W. Beecher, J.D. Cavalcoli, B.D. Athey, G.S. Omenn, C.F. Burant, H.V. Jagadish, Metscape: a Cytoscape plug-in for visualizing and interpreting metabolomic data in the context of human metabolic networks, Bioinformatics 26(7) (2010) 971-973.

[8] K.A. Brown, B. Chen, T.M. Guardado-Alvarez, Z. Lin, L. Hwang, S. Ayaz-Guner, S. Jin, Y. Ge, A photocleavable surfactant for top-down proteomics, Nature Methods 16(5) (2019) 417-420.

[9] F. Meier, A.-D. Brunner, S. Koch, H. Koch, M. Lubeck, M. Krause, N. Goedecke, J. Decker, T. Kosinski, M.A. Park, N. Bache, O. Hoerning, J. Cox, O. Räther, M. Mann, Online Parallel Accumulation–Serial Fragmentation (PASEF) with a Novel Trapped Ion Mobility Mass Spectrometer, Molecular & Cellular Proteomics 17(12) (2018) 2534-2545.

[10] N. Prianichnikov, H. Koch, S. Koch, M. Lubeck, R. Heilig, S. Brehmer, R. Fischer, J. Cox, MaxQuant Software for Ion Mobility Enhanced Shotgun Proteomics*, Molecular & Cellular Proteomics 19(6) (2020) 1058-1069.

[11] S. Tyanova, J. Cox, Perseus: A Bioinformatics Platform for Integrative Analysis of Proteomics Data in Cancer Research, in: L. von Stechow (Ed.), Cancer Systems Biology: Methods and Protocols, Springer New York, New York, NY, 2018, pp. 133-148.

[12] D.W. Huang, B.T. Sherman, R.A. Lempicki, Systematic and integrative analysis of large gene lists using DAVID bioinformatics resources, Nature Protocols 4(1) (2009) 44-57.

[13] J. Chong, O. Soufan, C. Li, I. Caraus, S. Li, G. Bourque, D.S. Wishart, J. Xia, MetaboAnalyst 4.0: towards more transparent and integrative metabolomics analysis, Nucleic Acids Research 46(W1) (2018) W486-W494.
